# IgG3 Hinge Architecture Expands Dengue Virus Breadth of A Quaternary Epitope Dependent Zika Virus Cross-Neutralizing Antibody

**DOI:** 10.64898/2026.03.17.712406

**Authors:** Sanjeev Kumar, Connor A. P. Scott, Lingling Xu, Kathryn Moore, Jacob V. Velden, Samuel Shih, Richard H. West, Sneh Lata Gupta, Naphak Modhiran, Mehul S. Suthar, Daniel Watterson, Jens Wrammert

## Abstract

Defining how antibodies achieve broad neutralization across antigenically related flaviviruses is essential for rational therapeutics and vaccine design. Here, we describe 3A06, a human monoclonal antibody that neutralizes dengue virus (DENV) serotypes 1-3 and zika virus (ZIKV). Notably, subclass switching to IgG3 exhibits expanded neutralizing activity against all four DENV serotypes. High-resolution cryo-electron microscopy (cryo-EM) analysis of 3A06 Fab bound to chimeric ZIKV virions shows that 3A06 engages a quaternary epitope spanning three E protomers. Functional analyses identify a major functional contribution from the light-chain, where somatic hypermutations are critical for neutralization potency, and reversion of improbable mutations substantially reduces neutralization and increases antibody dependent enhancement (ADE) at sub-neutralizing concentrations. Cryo-EM analyses of intact IgG3 in complex with chimeric DENV2 and DENV4 show that the conformational clexibility and extended IgG3 Fc-hinge enable bivalent engagement across E-protein rafts in multiple configurations, overcoming geometrical constraints that limit binding in the IgG1 format. Together, these results define a distinct mode of epitope recognition relative to previously characterized E-dimer epitope (EDE)-specific broadly neutralizing antibodies (bnAbs), and demonstrate how IgG3 subclass architecture can expand virus neutralization breadth. These insights inform the rational engineering of antibodies and vaccines targeting quaternary epitopes on flaviviruses to overcome viral diversity.

## INTRODUCTION

Flaviviruses such as dengue virus (DENV) and Zika virus (ZIKV) impose a significant global health burden, with over 400 million infections annually, yet effective strategies for achieving broad and safe vaccines or therapeutics remain limited ^1,2^. Antibody-mediated protection is central to vaccine and therapeutic strategies, but is constrained by serotype specificity and the potential risk of antibody-dependent enhancement (ADE) ^3,4^. The envelope (E) glycoprotein of flaviviruses, which mediates viral entry, is a major target of neutralizing antibodies (nAbs) ^3,5–7^. While monomeric E protein has often been used in vaccines and diagnostics, recent studies suggest that E protein dimers, which are present as quaternary structure on mature flavivirus virions, represent promising templates for rational vaccine design^3,4,8,9^.

The E protein of flaviviruses exists in different conformational states during the viral life cycle, including a trimeric state together with the virally encoded pr chaperone in immature viral particles and as E dimers which assemble into the rafts of the mature infectious virions ^10,11^. The majority of the existing neutralizing monoclonal antibodies (mAbs) recognize epitopes on the monomeric form of E protein targeting its fusion loop epitope (FLE) or EDIII domain ^12^. However, a few antibodies that selectively engage the so called envelope dimer epitopes (EDE) formed on the dimeric form of E protein, in the context of the pre-fusion conformation on mature virions, have recently been described ^5,6,13–18^. Antibodies capable of binding such quaternary epitopes have shown increased breadth and potency in recent studies, suggesting that targeting of dimeric E-protein conformation on the viral surface is a possible approach for designing broad flavivirus vaccines and therapeutics ^5,14,15^. The quaternary epitopes targeted on the E-protein are known to mediate broad DENV neutralization. However, it remains unclear how antibody intrinsic features, and IgG subclass architecture, govern epitope recognition, breadth, and enhancement risk.

We previously identified several DENV and ZIKV nAbs from the plasmablasts of DENV2 infected donors. One of those nAbs 33.3A06 (described as 3A06 throughout this study) showed potent ZIKV neutralization despite being isolated from a DENV2 infected donor ^7,12^. In this study, we carried out a detailed characterization of 3A06 and found that it recognizes a quaternary epitope spanning multiple E protein protomers, with no detectable binding to soluble E monomers. Cryo-electron microscopy analysis of the 3A06 Fab in complex with chimeric ZIKV particles confirmed this mode of recognition. Functional studies further showed that somatic hypermutations in the light chain make a major contribution to neutralization potency. We also found that antibody subclass influences neutralization breadth. When expressed as IgG3, 3A06 gained the ability to neutralize all four DENV serotypes, whereas the IgG1 form did not neutralize DENV4. Structural analysis of intact IgG3 in complex with DENV2 and DENV4 particles showed that the extended hinge region supports bivalent engagement of the virion surface, providing a structural basis for the broader activity observed in this subclass. Together, these findings provide insight into antibody recognition of flaviviruses and inform approaches for antibody design and vaccine development.

## RESULTS

### 3A06 antibody recognizes a quaternary E-protein dimer epitope

We previously reported the discovery of a fully human monoclonal antibody (mAb) 3A06 from the plasmablasts of a DENV2 infected donor^7,12^. The 3A06 mAb neutralized ZIKV more potently than DENV1-3 serotypes with up to 45-fold difference in potency, despite being originally isolated from a DENV2 infected donor. To understand the epitope specificity, we first assessed its binding profile using ELISA and biolayer interferometry. The 3A06 mAb binds to recombinant soluble E (sE) protein dimers captured on Streptactin-XT coated plates that were derived from DENV1-3 and ZIKV, but fail to engage DENV4 dimers. The antibody showed little or no detectable binding to sE monomers, the EDIII domain or to viral lysates (**Figures 1A and S1A-D**). Using a live virus-based focus reduction neutralization test (FRNT) assay, we observed that 3A06 neutralized DENV1-3 with FRNT_50_ values of 0.07 – 0.1 µg/ml. This is comparable to EDE1-C10 (0.3 – 0.6 µg/ml) and EDE2-A11 (0.6 µg/ml) mAbs, while modestly lower against ZIKV as compared to EDE1-C10 (**Figure 1B**). Biolayer interferometry (BLI) analysis revealed a heterogenous interaction of 3A06 in the micromolar (2.62 µM) to picomolar (1.78 pM) range against DENV and ZIKV E-protein dimers (**Figure 1C**). Together, these findings establish that 3A06 recognizes a quaternary epitope associated with multiple E-proteins.

**Figure 1:**
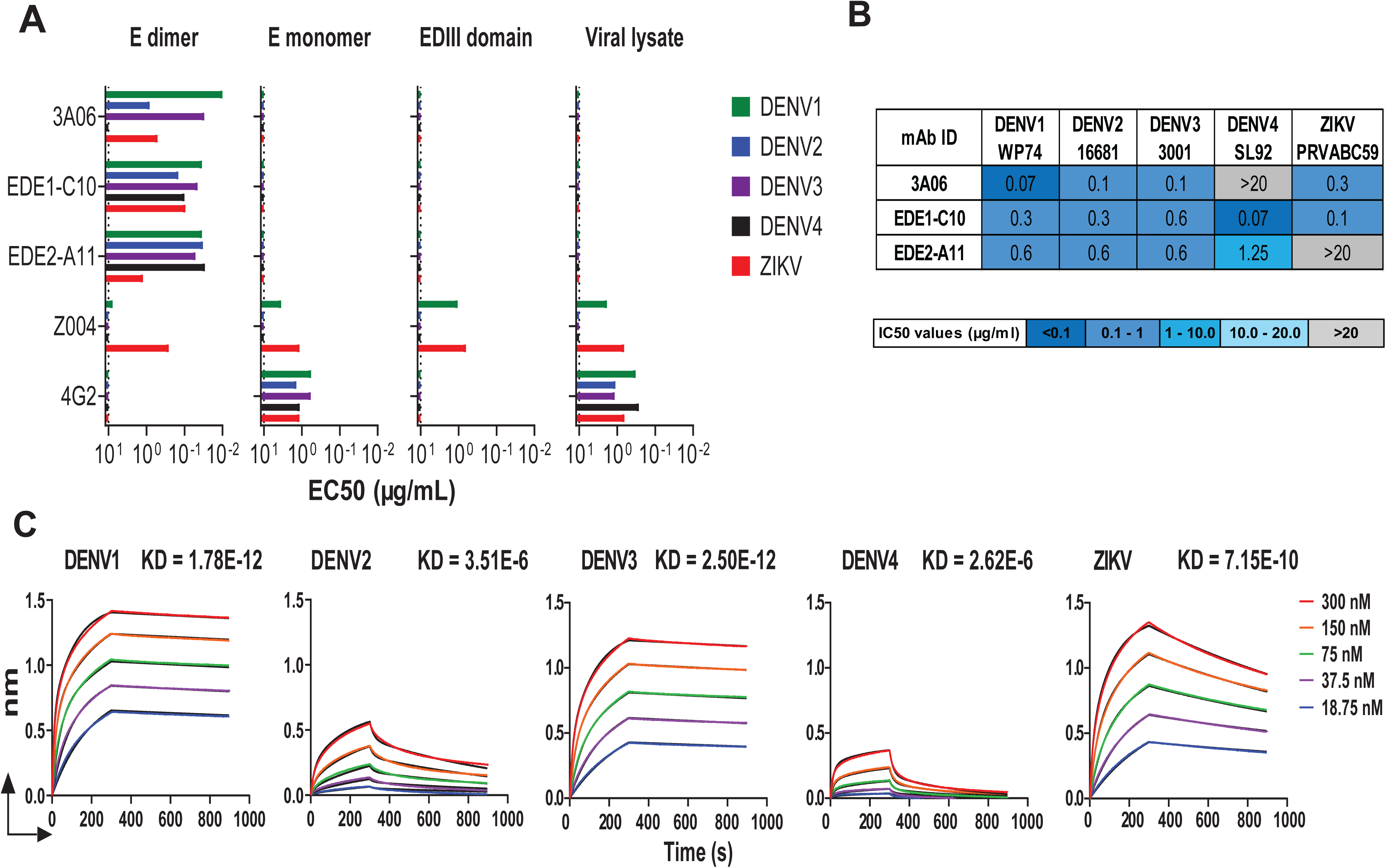
3A06 antibody selectively recognizes quaternary E-protein dimers and mediates broad and potent neutralization. **(A)** ELISA binding of 3A06 to recombinant E-protein dimers, monomers, EDIII, and viral lysates. The dotted line represents the limit of detection. Here, EDE1-C10 and EDE2-A11 antibodies were used as positive controls for E-dimers, FLE-specific antibody 4G2 was used here as control for E-monomers and viral lysates, and Z004 a ZIKV-EDIII specific antibody was used as control for EDIII domain for ZIKV. **(B)** Live-virus based focus reduction neutralization test (FRNT) assays data. **(C)** Biolayer interferometry (BLI) analysis reveals high-affinity binding to E dimers, with heterogeneous kinetics.

### Cryo-EM structure of 3A06 reveals recognition of a quaternary epitope spanning multiple E protomers

To understand the molecular basis of differences observed in 3A06 binding and neutralization against DENV and ZIKV, we first determined the structure of 3A06 Fab in complex with chimeric ZIKV virions. Here, we used the Binjari virus (BinJV) chimeric system ^19–21^, an insect-specific flavivirus, which has been shown to reproduce key structural and antigenic features of pathogenic flaviviruses while maintaining a high safety profile. The ZIKV structural proteins were presented on a Binjari virus backbone (herein bZIKV), to generate chimeric virions suitable for high-resolution cryo-EM analysis of 3A06 Fab binding. The 4.6 Å cryo-EM structure of bZIKV complexed with 3A06 Fab at 4°C showed that 3A06 binds a quaternary E-protein dependent epitope across three E protomers with 120 Fab fragments visible on the surface of the virion (**Figure 2A, Figure S2 and Table S1**). This structure was further refined using symmetry expansion of the asymmetric unit to produce a 3.1 Å map that could then be modelled (**Figure 2B**). The atomic model of bZIKV complexed with 3A06 Fab further highlighted the quaternary nature of the 3A06 epitope, showing extensive hydrogen bonding across three E protomers per Fab fragment (**Figure 2C**). Within the asymmetric unit, two 3A06 Fabs bind at the 2- and 3- fold axis respectively (**Figure 2C**). Each Fab binds across EDI and EDIII of one E protomer, and EDII of the neighboring two protomers, forming multiple bonds between the Fab and bZIKV (**Figure 3A, and 3B**). The VH of 3A06 buries 875 Å^2^ across two E protomers making six hydrogen bonds between its complementarity-determining region 2 (HCDR2) and HCDR3 loops and EDII of one E protomer and one hydrogen bond across to the neighboring EDI. R109 in the HCDR3 loop forms a salt bridge with D71 in the bZIKV bc loop, a highly conserved residue between ZIKV, and DENV serotypes 1-3 (**Figure 2C, 3B, 3C and Figures S3-S5**). Notably, DENV4 contains an alanine at position 71, which likely reduces the interaction strength of the 3A06 VH to DENV-4. The VL of 3A06 buries 673 Å^2^ with extensive hydrogen bonds between LCDR2 and EDIII and EDI of one E protomer, LCDR1 and the fusion loop and bc loop of the neighboring E protomer, and LCDR3 and the hi loop of the third E protomer (**Figure 2C, 3B and 3C**). Interestingly, several key hydrogen bonds for 3A06 to bZIKV were identified as low frequency mutations (2-10%) in germline mutation probability analysis (**Figure S6A, B**). S53, in the CDR-2 loop of 3A06 VL, which forms a hydrogen bond with K373 on EDIII of bZIKV E protomer A, and N33 which interacts with G104 of the E protomer C fusion loop. While G104 is highly conserved across flaviviruses, K373 is not conserved among DENV serotypes and ZIKV, with DENV1, 2, and 3 having a proline, and DENV4 having a valine at the equivalent position (**Figure 3B and Figures S3-S5**). Overall sequence conservation of the 3A06 epitope shows that the residues on EDII, including the FL, hi, and bc loop contacted by the VH and VL are more conserved between DENV1-4 and ZIKV compared to the quaternary contacts facilitated by the light chain primarily on EDI and EDIII (**Figure S3 and Figures S3-S5**). These contacts suggest that the VL interactions contribute substantially to the stabilization and recognition of the 3A06 epitope on ZIKV. Further, the structure provides insights into the quaternary epitope recognition by 3A06 which is distinct relative to previously reported quaternary epitope dependent DENV and ZIKV mAbs C10 ^22^, SIgN-3C ^23^, ZIKV-195 ^24^, and ADI30056 ^25^ (**Figure S7**). Together, these observations reveal structural features that distinguish the mode of interaction from previously characterized EDE-specific antibodies, such as the EDE1-C10 antibody.

**Figure 2:**
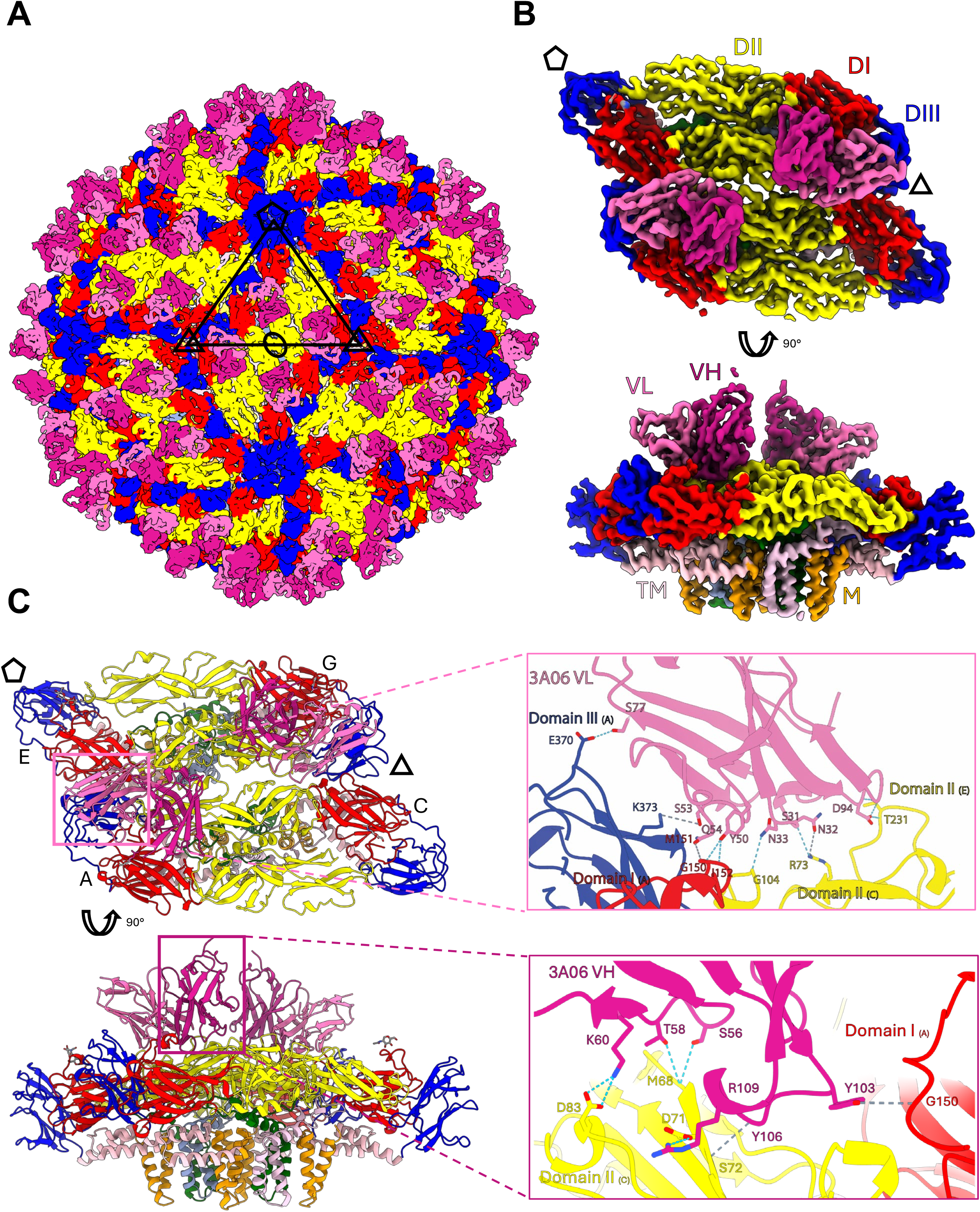
Cryo-EM structural analysis of 3A06 Fab in complex with ZIKV-BinJV. **(A)** 4.6 A{ cryo-EM reconstruction of bZIKV complexed with 3A06 Fab colored by E domains (EDI: Red, EDII: Yellow, EDIII: Blue) and Fab heavy variable (Magenta) and light variable (Pink) domains. The asymmetric unit of bZIKV is labelled with the ellipses, triangles and pentagon representing the 2-, 3- and 5-fold icosahedral axes. (**B)** 3.1 A{ symmetry expanded reconstruction of a pair of bZIKV E-M heterodimers complexed with 3A06 Fab. Density has been zoned to atomic model in panel C to display key contact sites for two fab fragments. **(C)** Top and side view of atomic model of bZIKV complexed with 3A06 Fab with insets highlighting key hydrogen bonds for 3A06 LV (pink) and 3A06 HV (magenta).

**Figure 3:**
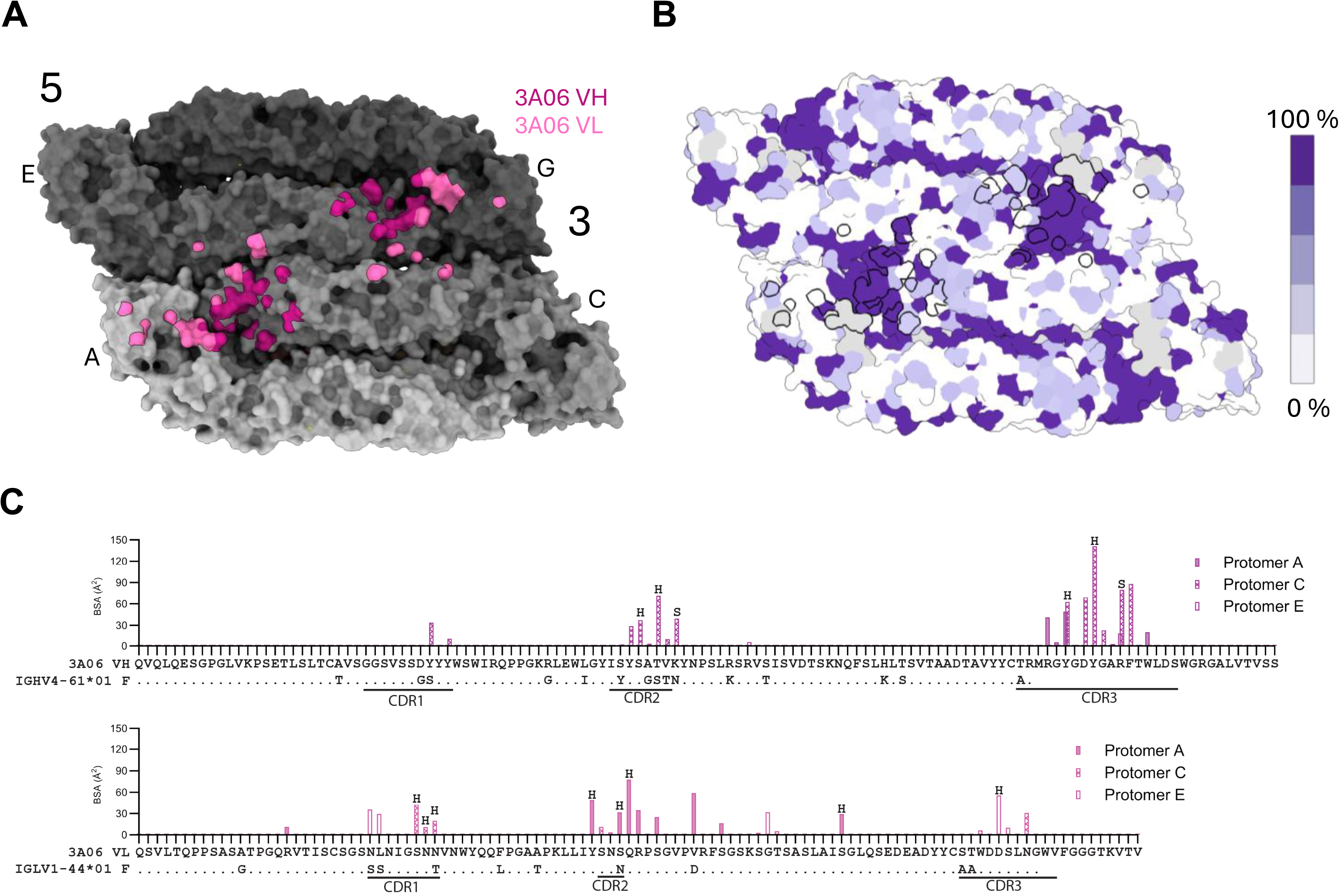
Binding footprint and conservation of the 3A06 Epitope. **(A)** Top view surface representation of 3A06 footprint on a pair of ZIKV E dimers with heavy chain variable region (HV) (magenta) and light chain variable region (LV) (pink). Each E monomer is labelled by chain (A, C, E, G) and with 5- and 3- fold symmetry axis. **(B)** The same model from **(A)** colored by sequence conservation of residues across DENV serotypes and ZIKV E. Coloring generated using Chimera X (v1.8). Footprint of 3A06 Fab shown in black outline. **(C)** Buried surface area plot showing which 3A06 residues are interacting with ZIKV E colored by E protomer with hydrogen (H) bonds labelled. Germline V-genes of both VH and VL are shown with the CDRs indicated.

### Somatic hypermutations in the 3A06 light chain strongly influence antigen recognition and neutralization

We further examined how somatic hypermutations (SHM) in the heavy (VH) and light (VL) chain variable regions contribute to neutralization and antigen recognition. To address this, we generated antibodies in which the VH region, the VL region, or both were reverted to their inferred germline sequences (iGL) as shown in (**Figure 4A**), allowing us to dissect the relative contributions of each chain to antigen recognition and neutralization potency. The iGLs were designed by reverting the VH or VL to their germline sequences while retaining the original CDRH3 and CDRL3 sequences based on sequence alignments generated by IMGT sequence analysis (**Figure S8**). Additionally, we evaluated the 3A06 antibody mutation probabilities using ARMADiLLO, which is an online server for identifying probable and improbable mutations ^26^. The improbable mutations (Imp) are low-probability amino acid changes determined by ARMADiLLO, which are amino acid substitutions in antibody variable regions that have a low probability (2% or lower) of occurring during the natural somatic hypermutation process, because they require multiple nucleotide changes ^26^. This analysis resulted in the identification of four improbable mutations in both the VH and VL genes ranging from 1-2% frequency in the 3A06 antibody (**Figure S6A-B**). Based on these analyses we generated additional combinations of 3A06 VH and/or VL chain genes with only the improbable mutation reverted to germline gene sequences (**Figure 4B**).

**Figure 4:**
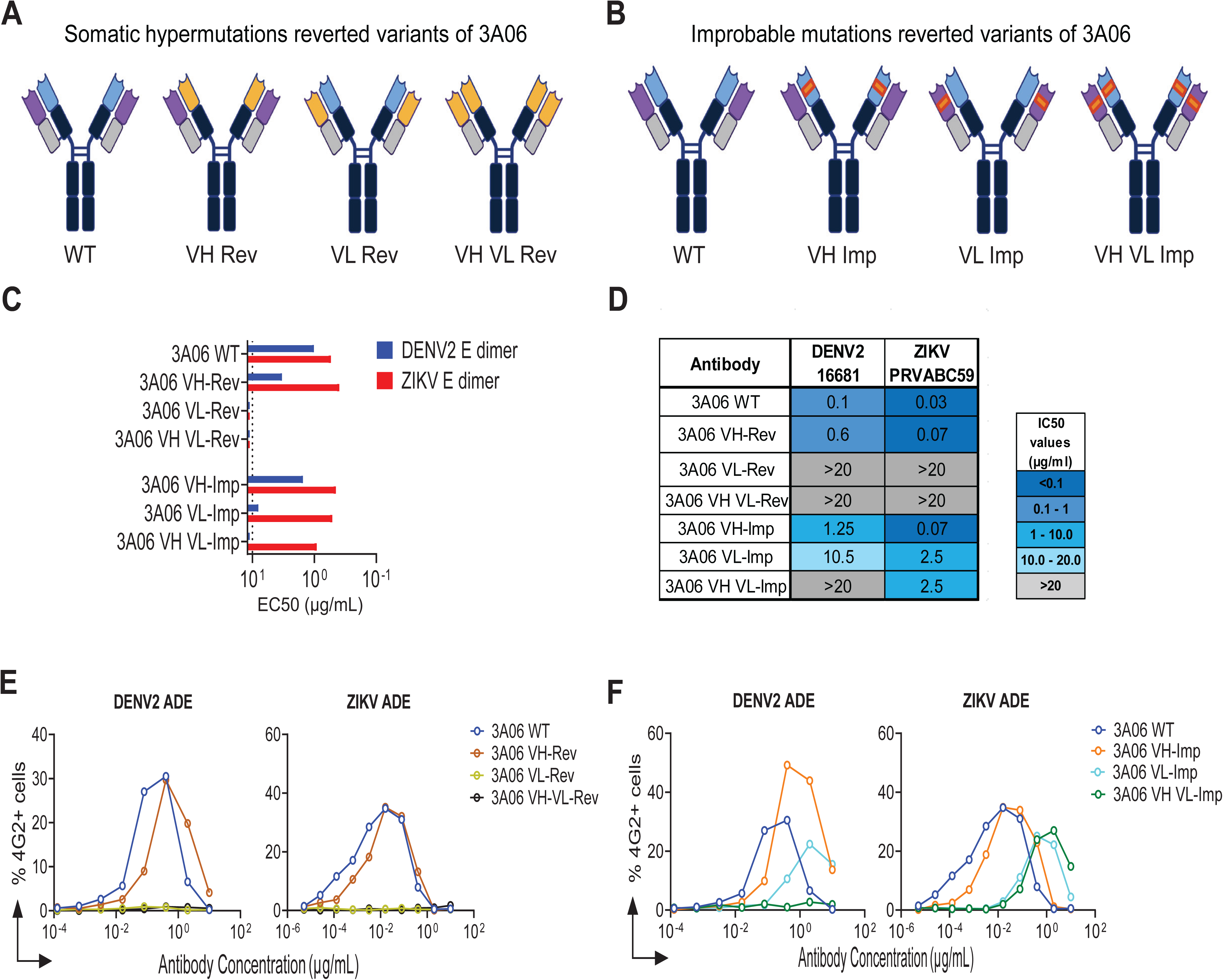
Neutralizing activity of 3A06 is primarily contributed by its light chain. **(A)** Pictorial presentation of 3A06 WT and its somatic hypermutation reverted iGL variants, variable heavy chain gene reverted VH-Rev, variable light chain gene reverted VL-Rev and both variable heavy and light chain genes reverted VH VL-Rev antibodies. **(B)** Pictorial of 3A06 WT and its improbable mutations reverted Imp variants, variable heavy chain gene reverted VH-Imp, variable light chain gene reverted VL-Imp and both variable heavy and light chain genes reverted VH VL-Imp antibodies. **(C)** ELISA binding of 3A06 iGL and Imp antibodies. The dotted line represents the limit of detection. **(D)** 3A06 iGL and Imp antibodies neutralization analysis by live-virus based FRNT assay. **(E)** In-vitro ADE analysis of 3A06 variants against DENV2 and ZIKV using U937 cells.

We found that antibodies based on the light chain variable region (iGL and Imp variants) are important for recognizing antigens. In comparison to the heavy chain-based variants, the light chain variants showed little or no binding to DENV2 and ZIKV E-protein dimers (**Figure 4C and S9**). The VH reverted 3A06 IgG1 antibody showed reduced neutralizing activity against both ZIKV and DENV2. However, the VL reverted and VH-VL reverted iGL 3A06 antibodies showed no neutralizing activity against ZIKV or DENV2 (**Figure 4D**). Similarly, the VH-Imp reverted 3A06 IgG1 antibody showed reduced neutralization potential against both ZIKV and DENV2. In comparison, the VL-Imp reverted antibody showed substantial reduction in neutralizing activity against DENV2 compared to wild-type 3A06 IgG1 antibody (**Figure 4D**). The VH VL-Imp reverted 3A06 antibody showed reduction in neutralization against ZIKV. Against DENV2, the VH VL-Imp reverted 3A06 antibody did not show any neutralization compared to the wild-type 3A06 IgG1 antibody (**Figure 4D**). Moreover, reversion of these key SHMs resulted in increased infection in U937 cells at sub-neutralizing concentrations, suggesting that affinity maturation modulate the balance between neutralization and enhancement in these assays (**Figure 4E and 4F**). These results demonstrate that SHMs in the 3A06 light chain variable gene make a major contribution to antigen recognition and neutralization potency against both DENV2 and ZIKV. Reversion of specific low-probability or improbable mutations markedly reduced neutralization potency, suggesting that affinity maturation of the light chain plays an important role in shaping the functional activity of 3A06 antibody.

### 3A06 is encoded by an uncommon combination of antibody variable genes

We next investigated the genetic basis of the structural and functional properties exhibited by 3A06. Sequence analysis of the variable regions revealed that 3A06 is encoded by a combination of heavy and light chain immunoglobulin genes not commonly observed among other flavivirus broadly neutralizing antibodies, including those targeting EDE1 (EDE1-C8, C10), EDE2 (EDE2-B7, A11), or other quaternary epitopes (SIgN-3C) (**Table 1 and Figure S8**). The heavy chain variable region is derived from IGHV4-61*01, IGHD4-17*01, and IGHJ5*01, while the light chain derived from IGLV1-44*01, IGHJ3*01 gene. The 3A06 heavy chain exhibits moderately high levels of somatic hypermutation (11.34%) compared to previously reported E-dimer specific mAbs. The light chain is somewhat less mutated (6.32%). 3A06 has a CDR3 length of 18 amino acids, and 11 amino acids in CDRL3 region, which are comparable to EDE1 specific antibodies EDE1-C10 and EDE1-C8 ^5^. The IGHV and IGLV gene combination seen in 3A06 may help explain why it recognizes its target differently from previously reported antibodies that bind quaternary epitopes. This also suggests that 3A06 arises from a gene pairing that is relatively uncommon.

**Table 1:**
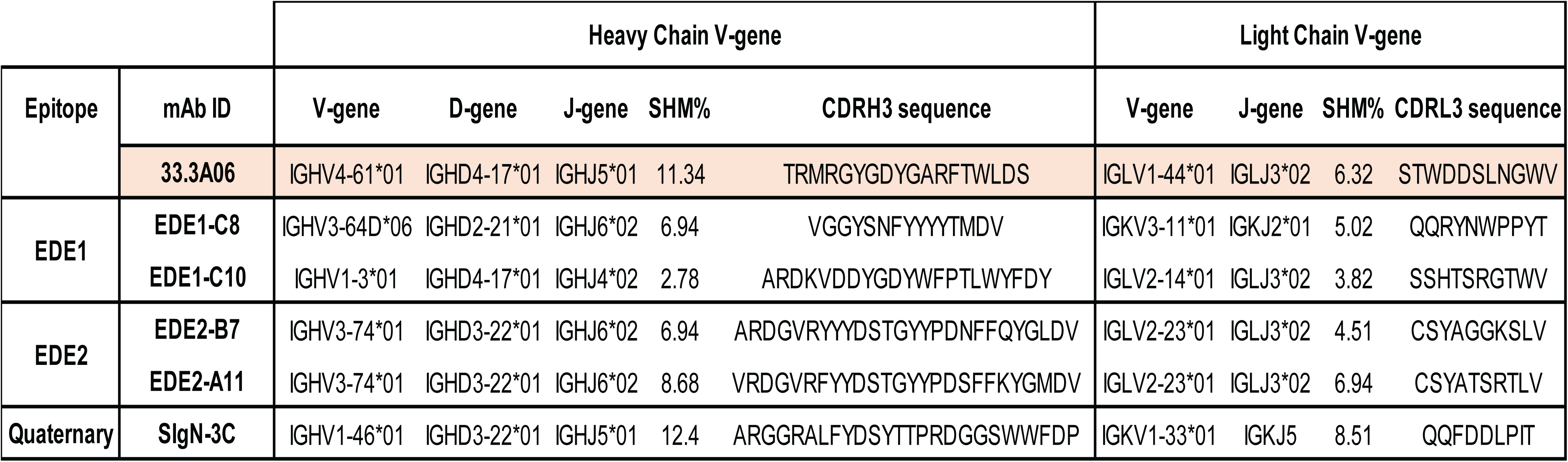
Comparison of immunogenetic features of quaternary E-protein epitope targeting antibodies to 3A06.

### 3A06 expressed in IgG3 subclass broadens neutralization against dengue and zika viruses

We next investigated whether antibody subclass influences the functional breadth of 3A06. While the IgG1 form effectively neutralizes DENV1-3 and ZIKV, it shows no detectable neutralization of DENV4. To determine whether Fc architecture could modulate this activity, we additionally generated IgG2, IgG3, and IgG4 variants and assessed their neutralization profiles across all four dengue serotypes (**Figure 5A**). In contrast to 3A06 IgG1, the IgG3 version showed higher binding to sE protein dimers of all four DENV serotypes and ZIKV (**Figure 5B and Figure S10**) and potent neutralization activity against DENV4 (**Figure 5C**), which was resistant to neutralization by 3A06 IgG1. Further, to assess the potential for 3A06 mediated ADE, we performed DENV2 and ZIKV infection assays using Fcγ receptor expressing U937 cells pre-incubated with varying concentrations of 3A06 in various Fc variants. The 3A06 IgG1 variant mediated moderate enhancement of DENV2 and ZIKV infection at sub-neutralizing concentrations (**Figure 5D**). In contrast, the IgG3 version induced higher levels of DENV2 and ZIKV ADE. The IgG2 subclass antibody didn’t mediate enhancement, consistent with its low affinity for Fcγ receptors ^27,28^.

**Figure 5:**
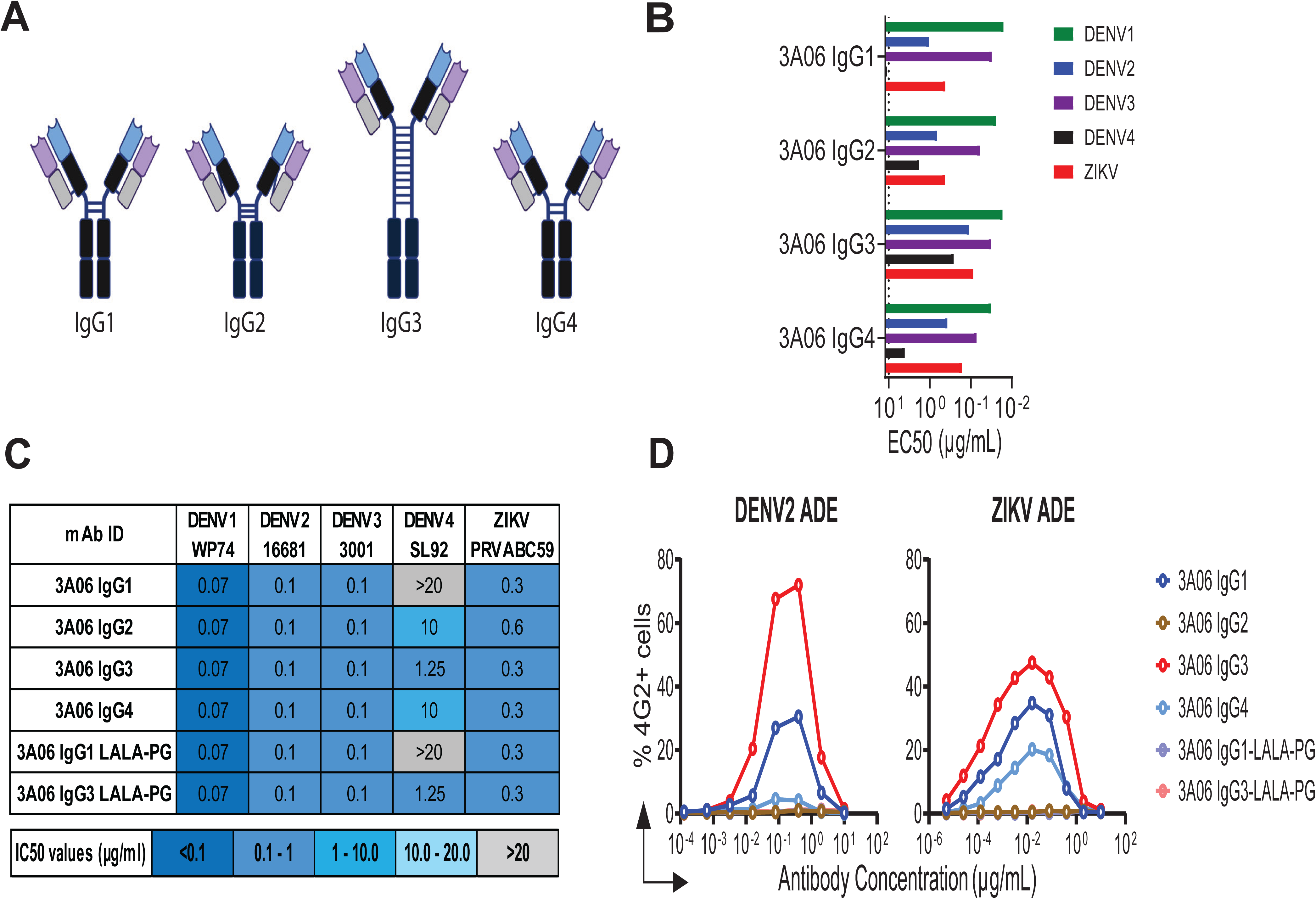
3A06 IgG3 antibody subclass version potently neutralizes DENV4. **(A)** Pictorial presentation of 3A06 IgG1, IgG2, IgG3 and IgG4 subclass antibodies. **(B)** ELISA binding of 3A06 IgG subclasses. The dotted line represents the limit of detection. **(C)** 3A06 neutralization analysis by live-virus based FRNT assay. **(D)** In-vitro ADE analysis of 3A06 IgG subclasses against DENV2 and ZIKV using U937 cells.

Based on these observations, we next engineered the L234A/L235A/P329G (LALA-PG) Fc variant of the 3A06 antibody in both IgG1 and IgG3 subclasses. The LALA-PG Fc mutant is well-known to abrogate Fcγ receptor binding in IgG1, and here we grafted the same mutations into IgG3 to determine whether these could also abolish Fcγ receptor binding in the IgG3 Fc backbone. We did not observe any change in neutralization activity between the LALA-PG IgG1 and IgG3 antibodies compared to their respective wild-type antibodies (**Figure 5C**). As expected, IgG1 LALA-PG antibody completely eliminated ADE. Interestingly, the same mutations incorporated into the IgG3 version were also effective in silencing ADE (**Figure 5D**). These findings indicate that antibody IgG3 subclass conversion influences the quaternary epitope accessibility, and functional neutralization breadth.

### 3A06 IgG3 hinge architecture facilitates bivalent virion engagement

In order to understand the unique capacity of 3A06 to bind and neutralize DENV4 when presented on an IgG3 backbone, we performed cryo-EM of whole IgG3 in complex with chimeric DENV2-BinJV and chimeric DENV4-BinJV and compared these to ZIKV-BinJV and DENV2-BinJV complexed with 3A06 Fab. Whole IgG3 format mAbs were purified by protein G and complexed with virions. Cryo-EM datasets were collected and complexes resolved by SPA. We determined cryo-EM structures of IgG3-bDENV2 at 4.4 Å resolution and IgG3-bDENV4 at 4.17 Å resolution. Comparing the resulting maps, the antibody density was the lowest for DENV2 complexed with 3A06 Fab, followed by DENV4 3A06 IgG3, DENV2 3A06 IgG3 and ZIKV 3A06 (**Figure 6A, Figure S2 and Table S2**), consistent with the observed binding levels. Similar to ZIKV, no antibody density was observed at the 5-fold proximal site, which results from a clash across the E rafts, and this configuration also abrogates the intra-dimer LC contacts critical for 3A06 binding. Although the occupancy was improved for the DENV2 bound to intact IgG3 compared with Fab alone, no clear Fc or hinge density was observed, most likely due to the inherent flexibility of these regions. Interestingly, extra density around the CH1 and CL domains was observed for DENV2 complexed with 3A06 IgG3 when compared to ZIKV and Fab (**Figure 6B, green**). In order to examine this in more detail, focused classification was carried out using symmetry expanded particle sets, focusing on either side of the CH1 and CL domains (**Figure 6B dashed circles**). This revealed two distinct classes, where the Fab densities at the 3-fold proximal site can adopt either a conformation with the CH1 and CL domains bent over towards the adjacent 2-fold position within the E raft (**Figure 6C, class 1**) or a conformation similar to the 3-fold site Fab (**Figure 6C, class 2**). These two conformations alter the relative positions of the CH1 linker region and therefore influence the potential for bivalent engagement of the full antibody molecule (**Figure 6D, Figure S11**). Notably, we observed Fab densities at the 2-fold position (intra-raft) in both classifications, while the antibody density at the adjacent 3-fold position (inter-raft) was stronger in the 2^nd^ class, consistent with the ability of the IgG3 to bind across the E rafts when in this more favorable CH1-CL conformation. This is different to what has been previously observed for DENV mAb C10 when complexed with virions as a F(ab)2’ ^22^, where only a single bivalent binding mode was observed with C10 adopting a conformation similar to our observed Class 1, but with no clear densities at the 2-fold sites. To examine this difference further, we investigated the relative distances of the first disulfide position of the hinge for C10 and 3A06 in IgG1 and IgG3 framework (**Figure S12**). We found that the hinge disulfide is least favorable for 3A06 in the IgG1 format, followed by C10. In contrast, in the IgG3 format, 3A06 readily retains the hinge disulfide while simultaneously engaging both 3-fold proximal epitopes within a raft. Based on these findings, the inability of 3A06 IgG1 to neutralize DENV4 is likely due to structural constraints that prevent effective multivalent binding to the virion. In contrast, the longer and more flexible hinge region of IgG3 enables it to span and engage multiple sites across the virion more effectively. This increased flexibility enhances binding strength or avidity and overall functional activity, thereby permitting neutralization. Overall, these results indicate that the extended hinge architecture of the IgG3 subclass facilitates bivalent engagement of the virion in multiple configurations, leading to expanded neutralization breadth.

**Figure 6:**
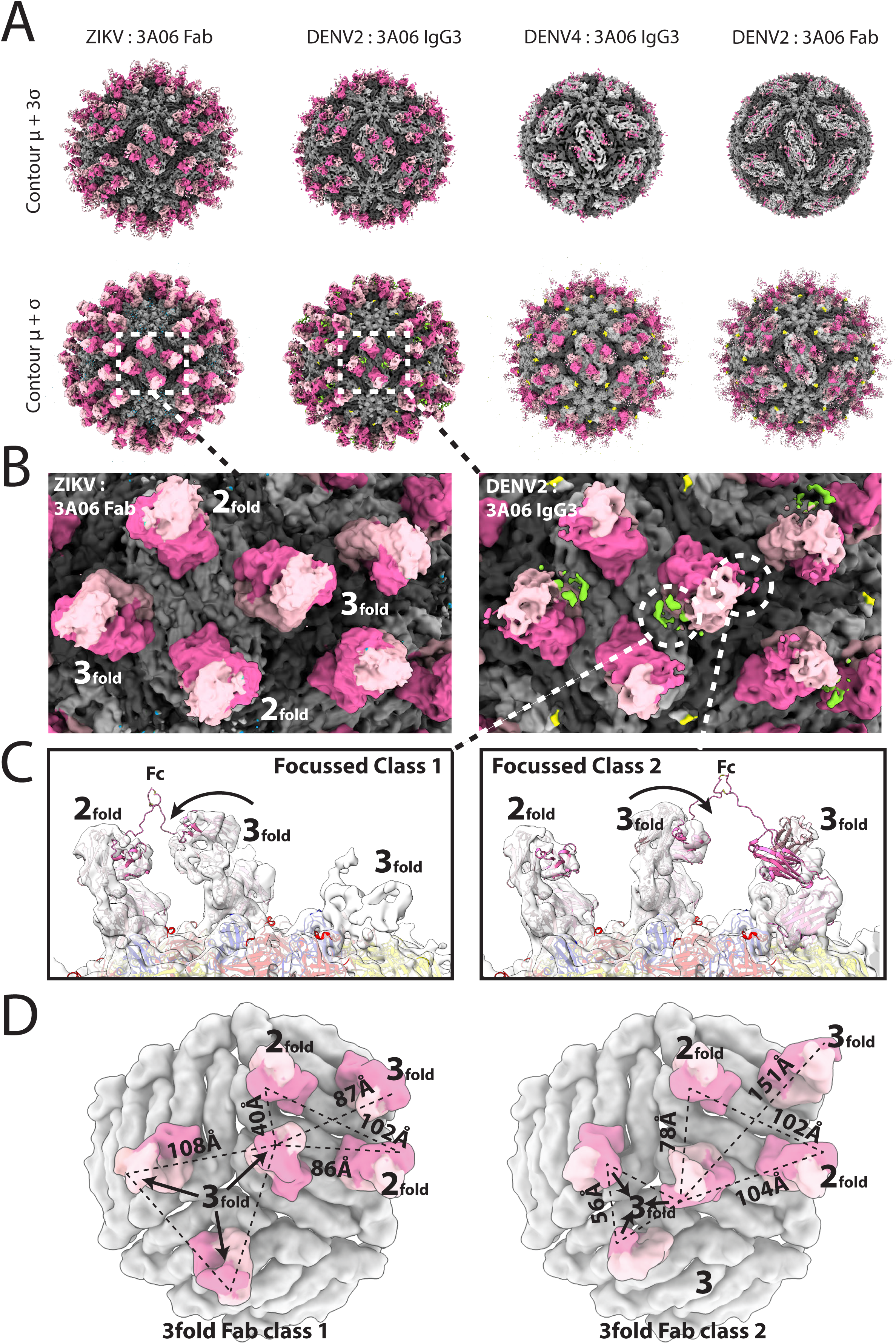
Cryo-EM structural analysis of 3A06 IgG3 complexed with DENV2 and DENV4. **(A)** Whole virion maps of bZIKV complexed with 3A06 Fab, bDENV2 complexed with intact 3A06 IgG3, bDENV-4 with intact 3A06 IgG3 and bDENV-2 complexed with 3A06 Fab. Virions colored in greys representing the 3 E proteins in the asymmetric unit, with the 3A06 HC in magenta and LC in pink. 2 different contour levels (mean+3σ top, mean+ σ below) are shown for each reconstruction to highlight the occupancy differences of 3A06 for each complex**. (B)** Close up view of the 3A06 densities shown for bZIKV:Fab (left) and bDENV2:IgG3 (right). Additional densities observed only in the DENV2 IgG3 bound map are colored yellow for the position of the N67 DENV glycan and green for an extra density adjacent to the 3-fold Fab constant domain. **(C)** Focused classification of the extra densities revealed two conformations for the-fold Fab on IgG3 bound DENV2, consistent with two alternate bivalent interactions provided by the IgG3 Fc linker reach. **(D)** Spatial arrangement of the Fab densities on the two conformations. Distances between the Glu216 (Eu numbering) residues in the HC CH1 are shown for all potential bivalent interactions that could be mediated by a single mAb.

## DISCUSSION

The comprehensive characterization of 3A06 as a quaternary E-protein conformation specific human monoclonal antibody with potent DENV and ZIKV cross-neutralizing capacity offers several conceptual advances. First, it advances the existing knowledge that heavy chain dominated paratopes primarily determine antigen specificity in anti-flavivirus antibodies. In the case of 3A06, we find that antigen binding, and neutralization potency are critically dependent on the light chain, as confirmed by functional assays based on chimeric constructs encoding iGL variants and improbable mutations reverted 3A06 antibody variants. Such light chain dependent interactions compared to previously described antibodies in which light chain contributions are more evenly distributed may open new avenues for epitope focused flavivirus vaccine design and antibody optimization strategies.

Second, our results show that IgG subclass plays an important role in shaping both neutralization breadth and antibody-dependent enhancement for 3A06. In the IgG1 format, 3A06 neutralizes DENV1-3 and ZIKV but does not neutralize DENV4, a serotype that is often less sensitive to broadly neutralizing antibodies. When expressed as IgG3, however, 3A06 gains the ability to neutralize all four DENV serotypes. This difference can be explained by structural features of the IgG3 hinge, which increase flexibility and allow more effective engagement of epitopes on the virion surface. Our cryo-EM analysis of intact 3A06 IgG3 in complex with DENV particles supports this interpretation. The data show that movement of the CH1 and CL domains at the 3-fold site allows the antibody to adopt more than one binding configuration, enabling engagement of both intra-raft and inter-raft epitopes. This behavior differs from what has been reported for the C10 antibody, where binding is limited to a single intra-raft configuration ^22^. These observations suggest that the IgG3 format enhances the ability of 3A06 to interact with the complex quaternary epitope present on DENV4. A central conclusion from this work is that the lack of DENV4 neutralization by 3A06 IgG1 is not due to a failure to recognize the epitope. Instead, it reflects an inability to meet the spatial requirements needed for effective binding.

The 3A06 epitope spans both 2-fold and 3-fold proximal regions, and neutralization requires bivalent engagement across E protein rafts. This creates a geometry-dependent requirement in which the antibody must bridge specific distances and orientations between adjacent sites. For DENV4, these constraints appear to be more restrictive. The shorter and less flexible hinge of IgG1 likely limits its ability to adopt conformations that support binding across rafts, resulting in predominantly monovalent interactions that are not sufficient for neutralization. In contrast, the longer hinge of IgG3 allows a wider range of Fab orientations, including those that support cross-raft binding. This enables stable bivalent engagement and increases overall binding strength, which allows neutralization to occur. Together, these findings define a subclass-dependent mechanism in which neutralization is influenced not only by epitope recognition but also by the physical properties of the antibody. More broadly, these results show that DENV4 is not escaping recognition by 3A06, but rather limiting the antibody’s ability to engage both binding sites at the same time. The IgG3 format overcomes this constraint by providing the flexibility needed for effective bivalent binding. In addition, we show that LALA-PG mutations effectively abrogate ADE in U937 cells in the IgG3 format, consistent with their established effects in IgG1 ^28–30^. This result indicates that Fc effector functions can be selectively minimized in IgG3 without disrupting its structural advantages for antigen engagement. Such control may be particularly important in the context of flaviviruses, where FcγR interactions contribute to ADE. Together, these findings support the feasibility of combining subclass-specific properties with Fc engineering to optimize antibody function.

The rare germline gene usage of 3A06 further supports the idea that some of the most effective broadly neutralizing antibodies arise from uncommon B cell lineages. The IGHV4-61/IGLV1-44 combination observed here is not reported in known DENV/ZIKV antibodies targeting the conformational epitopes present on the dimeric state of the E-protein. This highlights the possibility that natural DENV infection resulted in induction of E-dimer specific antibodies comprising different unbiased immunogenetic features. These findings highlight the need for expanded profiling of underrepresented B cell clones and non-public clonotypes. The quaternary epitope recognized by 3A06 is present only on intact virions and is not reproduced by monomeric E protein, suggesting that vaccine antigens must preserve native E-protein organization that is present on mature virions to elicit similar antibodies.

In summary, this study defines the structural and functional features of the human monoclonal antibody 3A06, which recognizes a quaternary epitope spanning three E protein protomers on the flavivirus surface. Structural analysis shows that 3A06 engages this interface through coordinated contacts involving both heavy and light chains. Functional data further demonstrate that mutations in the light chain make a substantial contribution to antigen recognition and neutralization potency. We also show that antibody subclass can influence antiviral activity. The IgG3 format, with its longer and more flexible hinge, supports bivalent engagement of adjacent epitopes on the virion surface and expands neutralization breadth, including activity against DENV4 that is not observed in the IgG1 form. These findings highlight how both paratope composition and antibody architecture shape recognition of quaternary epitopes on flavivirus particles. Therefore, our findings suggest that IgG3-based antibody formats may offer potential advantages when targeting structurally complex or spatially constrained epitopes on flaviviruses and other viruses, such as HIV, influenza virus, and SARS-CoV-2, where epitope accessibility is shaped by virion architecture. While these implications require further study, our findings support continued structural and functional evaluation of IgG3 antibodies across different viral systems. Together, this work provides a framework for understanding how antibody structure relates to function and may guide the design of vaccines and therapeutic antibodies with improved breadth and potency against flaviviruses and related pathogens.

## METHOD DETAILS

### Cell lines

Vero CCL-81 cells (ATCC), were maintained in Eagle’s Minimum Essential Medium (EMEM) (ATCC) medium supplemented with 10% fetal bovine serum (FBS) (HyClone) and antibiotic-antimycotic (1X) (Gibco). U937 cells (ATCC) were maintained in RPMI-1640 supplemented with 10% FBS, 2mM L-Glutamine (Gibco), 1mM 4-(2-hydroxyethyl)-1-piperazineethanesulfonic acid (HEPES) (Gibco), 1mM sodium pyruvate (Gibco), 1X Non-essential Amino Acids (NEAA) (Gibco) and 1X penicillin/streptomycin (Gibco). All these cells were cultured at 37°C with 5% CO_2_. The expi293^TM^ cells (Gibco) were maintained in expi293 expression medium (Gibco) and cultured at 37°C, 125 rpm shaking and 8% CO_2_.

### DENV and ZIKV ampli[ication and lysate preparation

DENV1 strain WestPac74, DENV2 strain 16681, DENV3 strain 3001, DENV4 strain Sri Lanka 92 and ZIKV strain PRVABC59 was obtained from the Centers for Disease Control and Prevention (CDC) and was passaged by infecting Vero CCL-81 cells at a multiplicity of infection (MOI) of 0.1 in EMEM (ATCC). After a 1 h infection at 37°C, the virus inoculum was removed and EMEM supplemented with 10% FBS and 1% antibiotic-antimycotic (1X) (Gibco) was added to the cells. Upon observation of severe cytopathic effect (CPE) 4 to 6 days (DENVs) and 48 hr for ZIKV, supernatants were collected and spun down at 4000 × g for 10 min at 4°C. Supernatant containing virus was supplemented with an additional 10% (vol/vol) FBS before freezing at –80 °C. Viruses were titred by focus forming assay on Vero CCL-81 cells before use.

The viral lysates were prepared as described previously ^7^. Briecly, to prepare all four DENV1-4 serotypes and ZIKV lysates, the remaining Vero CCL81 adherent cells and cell pellet from the virus-containing supernatant were washed twice with Dulbecco’s Phosphate Buffer Saline (DPBS) and then resuspended in the cell lysis buffer (10 mM Tris, 150 mM NaCl, 1% sodium deoxycholate, 1% Triton X-100, pH 7.4) supplemented with protease inhibitor (Thermo Fisher Scientific) and phosphatase inhibitor (Biovision). Mock lysate was prepared in a similar fashion with uninfected cells. Bradford assay was performed to quantitate total protein yield.

### Dimeric DENV and ZIKV Env antigen expression and puri[ication

All four serotypes DENV1-4, and ZIKV dimeric Env constructs were designed by incorporating dimer stabilizing disulfide bonds based on previous designs ^3^ with a c-terminal strep-tag II sequence were synthesized and cloned in pTwist Beta Glo Neo expression vector from Twist Biosciences. Transfections were performed as per the manufacturer’s protocol in expi293 cells (Thermo Fisher). Briecly, expi293 cells were seeded at a density of 2x10^6^ cells/ml in Expi293 expression medium and incubated at 37°C and 125 rpm with 8% CO_2_ overnight. The next day, 2.5x10^6^ cells/ml were transfected using ExpiFectamine^TM^ 293 transfection reagent (ThermoFisher). Next morning, enhancers 1 and 2 were added along with 0.5% of antibiotic-antimycotic solution (Gibco). The cells continued to grow for 5 days at 37°C, 125 rpm, 8% CO_2_ incubator. The cells were removed by centrifugation at 6,000g for 20 minutes at room temperature and dimeric env protein-containing supernatant was collected. The supernatant was filtered through 0.45 um filter assembly (VWR) and loaded onto pre-washed Strep-Tactin®XT 4Flow® resin (IBA Lifesciences) for affinity purification. Resin washed with 1x wash Buffer (buffer W) containing 25mM Tris, 150mM NaCl and 1mM EDTA with pH 8.0 followed by Env protein elution in elution buffer 1x BXT buffer (IBA). Eluted protein dialyzed against DPBS and concentrated. The concentrated protein ran onto a Superdex200 Increase 10/300 column and protein eluted as dimeric Env collected.

### Monoclonal Antibody Expression and Puri[ication

Human mAbs used in study were generated as previously described ^7,12^. First, 3A06, IgG subclasses variants (IgG1, IgG2, IgG3 and IgG4) and IgG1 Fc variants, L234A/L235A/P329G (LALA-PG) ^29,30^, were generated by replacing the constant region of the wild-type IgG1 heavy chain expression vector with synthesized constructs (Twist Bioscience). The variable heavy and light chain sequences of control antibodies EDE1-C10 ^16^, and EDE2-A11 ^5^ were synthesized (Twist Bioscience) and cloned into human IgG1 expression vectors based on PDB IDs 7A3N and 5N0A respectively. Next, the variable heavy and light chain sequences encoding expression plasmids were transiently expressed in expi293F cells for 5 days and secreted antibodies were purified from supernatants using protein A coupled sepharose beads (Pierce) as per manufacturer’s instructions.

### ELISA assays

The binding potential of mAbs to ZIKV and DENV E protein antigens (Monomer, EDIII domain) and viral lysates was determined by ELISA. Briecly, the recombinant DENV1-4, ZIKV antigens and viral lysates were coated in separate Nunc Maxisorp ELISA plates (Thermo Scientific) at a concentration of 1 μg/ml in DPBS overnight at 4°C. Next morning, the plates were washed thrice with DPBS containing 0.05% Tween-20 (DPBST) and blocked with DPBST containing 1% BSA (ELISA buffer) for 1.5 hr at room temperature. Then, mAbs with serial 3-fold dilution in ELISA buffer, starting at 10 μg/ml concentration were added to the plates and incubated at room temperature for 1.5 h. The plates were washed thrice with DPBST, and the IgG signals were detected by incubating with peroxidase-conjugated anti-human IgG (Jackson ImmunoResearch) for 1 h. The plates were washed thrice with DPBST and developed using an o-phenylenediamine substrate (Sigma) in 0.05M phosphate-citrate buffer (Sigma Aldrich) containing 0.012% hydrogen peroxide. The absorbance values were measured at O.D. 490 nm. The absorbance values were plotted against the concentration of antibodies.

DENV and ZIKV E-protein dimer-based ELISA assays were performed using StrepTactinXT coated plates. StrepTactinXT coated microplates (IBA lifesciences) do not require any functionalization or blocking steps prior to protein immobilization. Briefly, 100 µL of Strep-Tagged purified E-protein dimers in ELISA buffer at 1 µg/mL concentration were dispensed in the corresponding wells for protein immobilization by a 2 h incubation at room temperature. Following a triple wash with DPBST to remove unbound proteins, three-fold serial dilutions of test mAbs and control mAbs (EDE1-C10, EDE2-A11, Z004 and 4G2) starting at 10 μg/ml concentration in ELISA buffer were added and incubated for 1.5 h at room temperature. After 3 washes with DPBST, peroxidase-conjugated anti-human IgG (Jackson Immunoresearch) diluted 1:3000 in ELISA buffer was added and incubated for 1 h, followed by three washes with DPBST. Plates were developed with o-phenylenediamine substrate (Sigma-Aldrich) in 0.05 M phosphate-citrate buffer (Sigma-Aldrich) pH 5.0, containing 0.012% hydrogen peroxide (Thermo Fisher Scientific). Absorbance was measured at 490 nm to obtain the binding data.

### Octet BLI analysis

Octet biolayer interferometry (BLI) was performed using an Octet Red96 instrument (ForteBio, Inc.). Briecly, mAbs were captured on ProA biosensors at 5 μg/ml concentration in BLI running buffer (0.1% BSA, 0.001% Tween20 in 1x DPBS). Binding kinetics were tested with serial 2-fold serially diluted DENV1-4 and ZIKV E-protein dimers (100 nM to 6.25 nM). The baseline was obtained by measurements taken for 120 s in BLI running buffer. The biosensors were dipped to obtain association phase for 300 s in wells containing serial dilutions of E-protein dimers. Then, the sensors were immersed in BLI running buffer for as long as 600 s to measure the dissociation phase. Sensors were regenerated in glycine pH 2.0 buffer and immediately neutralized by dipping in Tris pH 9.0 buffer. The mean K_on_, K_off_ and apparent KD values of the mAbs binding affinities for E-protein dimers were calculated from all the binding curves based on their global cit to a 1:1 Langmuir binding model using Octet software version 12.0.

### Preparation of 4G2 Antibody

Pan-clavivirus anti-envelope mAb 4G2 was isolated from the supernatant of mouse hybridoma (D1-4G2-4–15; ATCC) grown in hybridoma medium (Thermofisher Scientific), and purified using protein G beads (Pierce) according to the manufacturer’s recommendations. The antibodies were stored in 1x PBS for in-vitro cell culture assays and with 0.05% sodium azide at 4°C for the binding assays.

### Live virus-based focus reduction neutralization test (FRNT) assays

The neutralization potential of mAbs and serum samples was determined by FRNT as described previously ^7^ with a few modifications. Two-fold serially diluted mAbs starting at 20 μg/ml concentration were incubated with a previously titrated amount of virus (∼100 focus forming units) of ZIKV or DENV1-4 for 1 h at 37 °C. Vero CCL81 cell monolayers in 96-well plates were subsequently infected with the mixture for 1 h at 37 °C. An overlay containing 1% (wt/vol) methylcellulose (Sigma) was added to the cells. After 48 h (ZIKV) and 72 h (DENV) incubation at 37 °C, the cells were washed and fixed with a 1:1 mixture of acetone and methanol at room temperature. Foci were stained using 4G2 at 0.5 μg/ml concentration for 2 h at room temperature followed by HRP-linked anti-mouse IgG (Cell Signaling) for 1 h and developed using TrueBlue Peroxidase substrate (KPL) for 2 h at room temperature. At each step, plates were washed thrice with 1x DPBS and Foci were imaged using a CTL-Immunospot S6 Micro Analyzer.

### Antibody Dependent Enhancement (ADE) assay

ADE assay was performed as described previously ^7^. Briecly, serially diluted mAbs were incubated with ZIKV with an MOI of 0.5 for 1 h at 37 °C. The immune complexes were added in a 96-well plate containing 2x10^4^ U937 cells (ATCC) per well in RPMI-1640 with 10% (vol/vol) FBS. The Cells were infected for 24 h at 37 °C. The cells were then washed and fixed/permeabilized using BD intracellular staining reagents Fix/Perm Solution (BD biosciences) and Perm/Wash Buffer (BD biosciences) according to the manufacturer’s protocol. Cells were stained with in-house prepared 4G2 antibody conjugated to Alexa cluor 594 for 1h for 25 min. The percentage of infected (4G2^+^) cells was determined using clow cytometry using the Cytek Aurora clow cytometer followed by analysis using FlowJo software v10.

### Virus and antibody puri[ication for Cryo-EM

Chimeric viruses containing the non-structural genes of Binjari virus (BinJV) and prME genes of ZIKV (Natal, Genbank: NC_035889)), DENV-2 (ET300, Genbank: EF440433), and DENV-4 (ET288, Genbank: EF440435) were generated using circular polymerase extension reaction as previously described ^19,20,31^. To ensure mature virion preparations the prM furin cleavage site of DENV-2 and DENV-4 prME was modified to an Incluenza H5 HA furin cleavage site (RERRRKKRS). Virus recovered from transfected C6/36 cells were passaged to P1 and propagated in C6/36 cells (MOI: 0.1). Clarified viral supernatant was then purified as previously described with PEG precipitation, followed by a 20% sucrose cushion, and 25-40% potassium tartrate gradient ^21,32^. Gradient fractions containing virus were then harvested and used for Cryo-EM imaging. Fab fragments of 3A06 were generated by cloning the HV region of 3A06 into a mammalian expression vector containing a C-terminal C-Tag (EPEA) ^33,34^. Plasmids encoding the HV Fab and LC were then transfected into ExpiCHO cells, incubated for 7 days at 37 °C shaking, with the clarified supernatant then purified using an CaptureSelect™ C-tagXL Pre-packed Column on an AKTA-Pure FPLC.

### Cryo-electron microscopy

Chimeric clavivirus were complexed with excess 3A06 Fab fragments or IgG3 for 1 hour at 4 °C before being applied to glow discharged R3.5/1 holey carbon grids (Quantifoil). Grids were then vitrified using an EM GP2 (Leica Microsystems) set at 4 °C and 95% relative humidity, grids were pre-blotted for 10 secs, followed by blotting for 3 secs and immediately plunge-frozen in liquid ethane. Grids were transferred under liquid nitrogen either to a CRYO ARM^TM^ 300 (JEOL JEM-Z300FSC) transmission electron microscope operated at 300 kV, equipped with a K3 direct electron detector (Gatan) and an omega in-column energy filter operated at 20 eV (JEOL), or to a CRYOARM^TM^ 200 TEM operated at 200 kV, equipped with a K2 Summit direct electron detector (Gatan) and an omega in-column energy filter operated as above. Movies on both microscopes were recorded using SerialEM at 60,000 x magnification ^35^.

### Cryo-EM processing, single particle analysis, and model building

Cryo-EM datasets were motion corrected, and the contrast transfer function (CTF) parameters were estimated using *cis*TEM ^36^. Virus-Ab particles were automatically picked, extracted, and classified over multiple rounds of 2D classification. The initial 3D reconstructions on *cis*TEM were caried out ab initio, followed by 3D refinement using the low-resolution ab into model as a reference. High-resolution asymmetric focused reconstructions were carried out by symmetry expansion using *cis*TEM as previously described ^21,37^. Each image was subtracted 60 times using each of the 60 icosahedral ASUs generating a dataset of isolated asymmetric units which were automatically centred. The dataset of ASUs was then further refined using *cis*TEM without symmetry applied. The initial atomic model of bZIKV complexed with 3A06 Fab was generated using an existing cryoEM structure of ZIKV virus (PDB: 6CO8) as a template with an AlphFold3 generated model of 3A06 Fab. All modelling was performed using iterative cycles of ISOLDE (v. 1.2.1), COOT (v.0.8.9.2), and PHENIX (v.1.19.2) ^38–40^. The geometry and quality of the model were evaluated through a combination of PHENIX and the wwPDB Validation system. For focussed classification of bDENV-2 complexed with 3A06 IgG3, independent focussed classification of the 3-fold proximal Fab fragment was performed by masking the density either side of the 3-fold fab on the ASU followed by 3D classification using Relion 5.0.1. Particle stacks of Relion classes were then reimported into cisTEM and manual 3D refinement was performed to generate final focus classified ASU maps.

### Immunogenetic analyses of antibody genes and envelope protein sequence alignment

The immunogenetic analysis of both heavy chain and light chain germline assignment, framework region annotation, determination of somatic hypermutation (SHM) levels (nucleotides) and CDR loop lengths (amino acids) was performed with the aid of IMGT/HighV-QUEST (www.imgt.org/HighV-QUEST) ^41,42^. The envelope protein sequence alignments were done using the MAFFT version 7, a multiple alignment program for amino acid or nucleotide sequences (https://mafft.cbrc.jp/alignment/server/index.html) ^43,44^.

### Statistical analysis

Graphpad Prism version 9.0 was used for all statistical analyses.

## DATA AND MATERIALS AVAILABILITY

Atomic coordinates and Cryo-EM maps for reported structures are deposited into the Protein Data Bank (PDB) and the Electron Microscopy Data Bank (EMDB) with accession codes PDB: 24ZZ and EMDB: 69949 for chimeric virus between Binjari virus and zika virus in complex with 3A06 Fab; EMDB: 69863 for chimeric virus between Binjari virus and dengue virus serotype-2 (ET300) in complex with 3A06 Fab; EMDB: 69785 for chimeric virus between Binjari virus and dengue virus serotype-2 (ET300) in complex with 33.3A06 immunoglobulin G3; EMDB: 69800 for chimeric virus between Binjari virus and dengue virus serotype-4 in complex with 33.3A06 immunoglobulin G3. This paper does not report original code. All data needed to evaluate the conclusions in the paper are present in the paper and/or the supplemental information. Any material request or additional information required to reanalyze the data reported in this paper is available from the corresponding author upon request.

## ACKNOWLEDGEMENTS

We thank and acknowledge the facilities, and the scientific and technical assistance of the Australian Microscopy & Microanalysis Research Facility at the Centre for Microscopy and Microanalysis, The University of Queensland.

## FUNDING

This work was funded in part by NIH/National Institute of Allergy and Infectious Diseases (NIAID) Grants R01AI149486 (to J.W. and M.S.S.), R56AI110516 (to M.S.S.) and DBT/Wellcome Trust India Alliance Early Career Fellowship grant IA/E/18/1/504307 (to S.K.).

## AUTHOR CONTRIBUTIONS

Conceptualization, S.K., D.W., and J.W.; methodology, S.K., N.M., M.S.S., D.W., J.W.; investigation, S.K., C.A.P.S., L.X., K.M., J.V.V., S.S., R.H.W., S.L.G., N.M.; visualization, N.M., M.S.S., D.W., J.W.; funding acquisition, S.K., M.S.S., D.W., J.W.; project administration, S.K., L.X., R.H.W.; supervision, M.S.S., D.W., J.W.; writing – original draft, S.K., C.A.P.S., M.S.S., D.W. and J.W.; writing – review & editing, all authors.

## DECLARATION OF INTERESTS

All other authors declare no competing interests.

## Notes

### Competing Interest Statement

The authors have declared no competing interest.

### Summary of Updates

This version of the manuscript has been substantially revised from a preliminary draft previously posted on bioRxiv for internal author feedback and grant-related dissemination purposes. The revisions include updated and finalized main figures, additional data analysis, improved methodological clarity, and expanded interpretation of results. Structural data have now been deposited in the Protein Data Bank (PDB) and EMDB, and associated accession details have been updated in the manuscript. The text has been edited for clarity, coherence, and completeness throughout. No version of this manuscript has been submitted to or reviewed by any journal prior to this submission.

