## Supplementary File for "IgG3 Hinge Architecture Expands Dengue Virus Breadth of A Quaternary Epitope Dependent Zika Virus Cross-Neutralizing Antibody"

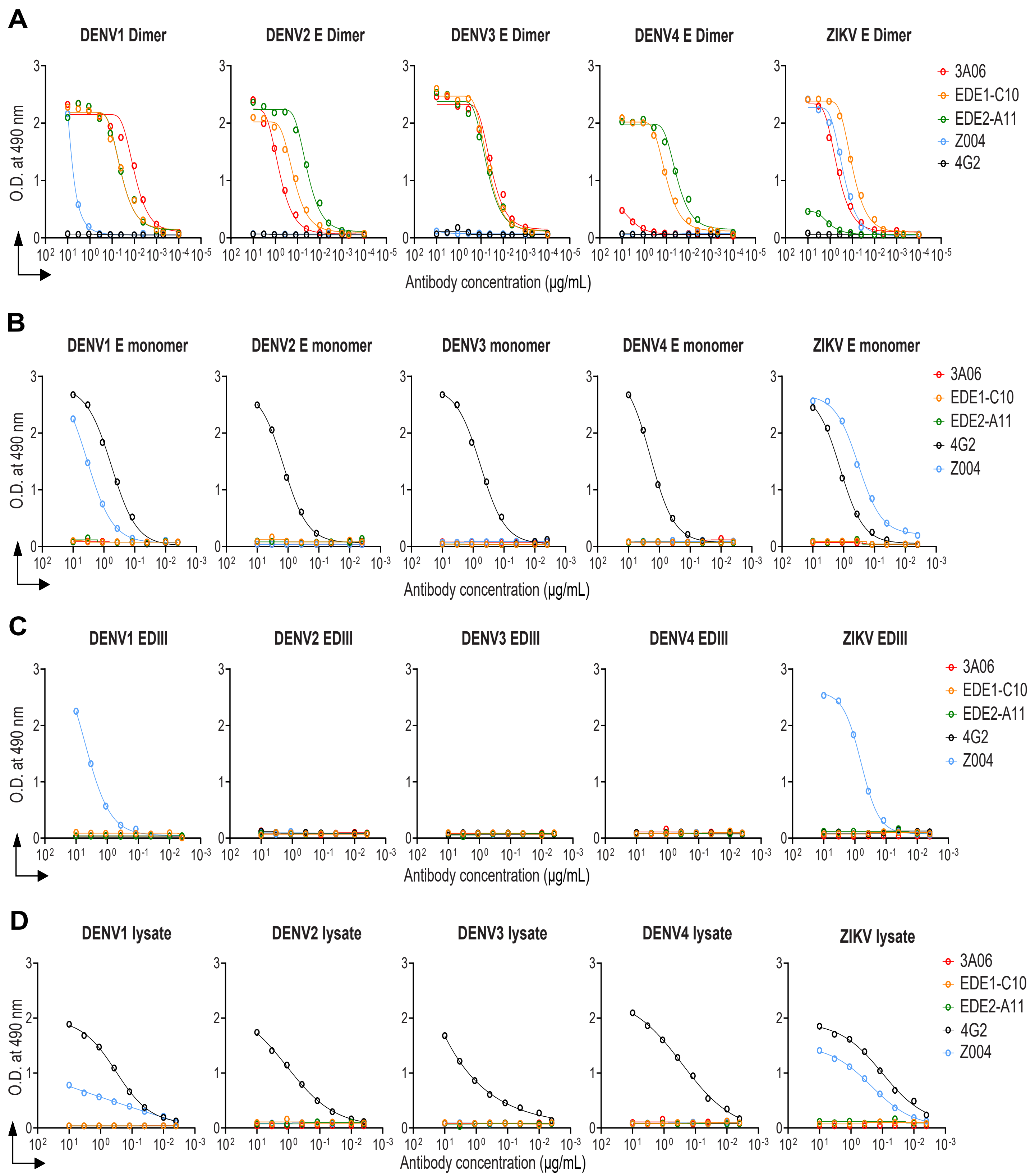


**Figure S1: 3A06 antibody binding analysis to DENV1-4 and ZIKV antigens.** Shown are the **(A)** E-protein dimer specific binding data. **(B)** sE monomers binding data. **(C)** EDIII domain binding data and **(D)** viral lysates binding data. Here, E-protein quaternary epitope dependent mAbs EDE1-C0, and EDE2-A11 were used as positive controls. A fusion loop epitope (FLE) specific mAb 4G2 was used as a negative control for FLE closed E-dimers and EDIII domains, positive control for sE monomers. ZIKV EDIII specific mAb Z004 was used here a positive control for ZIKV antigens, that also showed cross-reactivity to DENV1 EDIII domain.

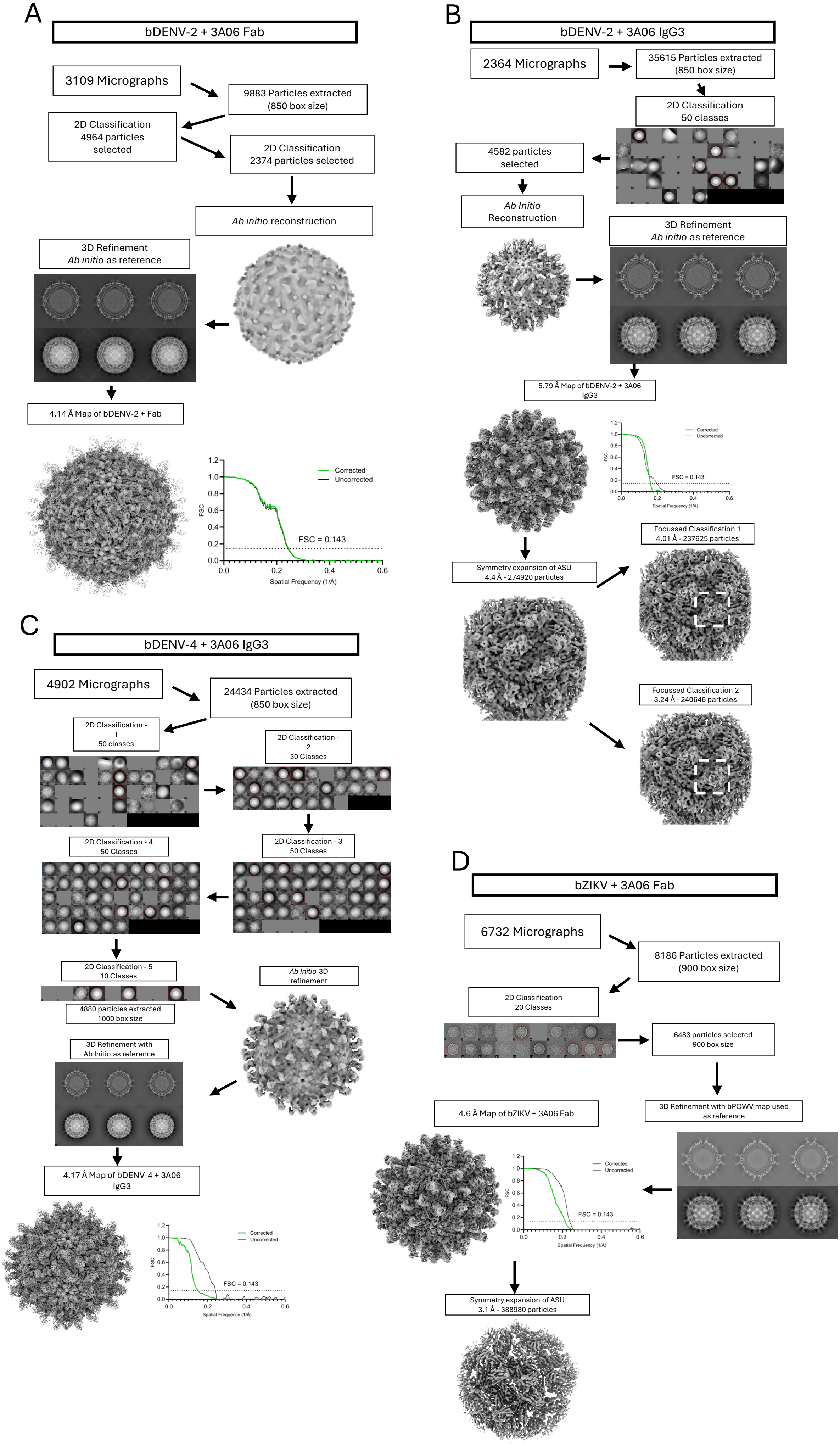

A
bDENV-2 + 3A06 Fab
3109 Micrographs
9883 Particles extracted (850 box size)
2D Classification 4964 particles selected
2D Classification 2374 particles selected
Ab initio reconstruction
3D Refinement Ab initio as reference
4.14 Å Map of bDENV-2 + Fab
Corrected
Uncorrected
FSC = 0.143
FSC
Spatial Frequency (1/Å)
B
bDENV-2 + 3A06 IgG3
2364 Micrographs
35615 Particles extracted (850 box size)
2D Classification 50 classes
4582 particles selected
Ab Initio Reconstruction
3D Refinement Ab initio as reference
5.79 Å Map of bDENV-2 + 3A06 IgG3
Symmetry expansion of ASU 4.4 Å - 274920 particles
Focussed Classification 1 4.01 Å - 237625 particles
Focussed Classification 2 3.24 Å - 240646 particles
C
bDENV-4 + 3A06 IgG3
4902 Micrographs
24434 Particles extracted (850 box size)
2D Classification - 1 50 classes
2D Classification - 2 30 Classes
2D Classification - 3 50 Classes
2D Classification - 4 50 Classes
2D Classification - 5 10 Classes
4880 particles extracted 1000 box size
Ab Initio 3D refinement
3D Refinement with Ab Initio as reference
4.17 Å Map of bDENV-4 + 3A06 IgG3
D
bZIKV + 3A06 Fab
6732 Micrographs
8186 Particles extracted (900 box size)
2D Classification 20 Classes
6483 particles selected 900 box size
3D Refinement with bPOWV map used as reference
4.6 Å Map of bZIKV + 3A06 Fab
Symmetry expansion of ASU 3.1 Å - 388980 particles

**Figure S2: CryoEM workflows for data presented in Figure 2, Figure 5 and Figure S7.** Motion correction and CTF estimation were performed on all datasets prior to particle picking. **(A)** Workflow of bDENV-2 complexed with 3A06 Fab displaying initial micrograph number, initial particle picks, multiple rounds of 2D classification and *ab initio* reconstruction of virion map. *Ab initio* model was then used for 3D refinement resulting in a 4.14 Å (0.143 FSC cut-off, shown next to final map) cryoEM map of bDENV-2 complexed with 3A06 Fab. **(B)** Workflow of bDENV-2 complexed with 3A06 IgG3 displaying initial micrograph number, initial particle picks, 2D classification and *ab initio* reconstruction of virion map. *Ab initio* model was then used for 3D refinement resulting in a 5.79 Å (0.143 FSC cut-off, shown next to final map) cryoEM map of bDENV-2 complexed with 3A06 IgG3. Virion map was further refined using symmetry expansion with no symmetry (C1) applied. Independent focussed classification of the 3-fold proximal Fab fragment was performed by masking the density either side of the 3-fold fab followed by 3D classification using Relion 5.0.1. Particle stacks of Relion classes were then reimported into cisTEM and manual 3D refinement was performed to generate final focus classified ASU maps of two distinct classes of 3-Fold fab position. **(C)** Workflow of bDENV-4 complexed with 3A06 IgG3 displaying initial micrograph number, initial particle picks, multiple rounds of 2D classification and *ab initio* reconstruction of virion map. *Ab initio* model was then used for 3D refinement resulting in a 4.17 Å (0.143 FSC cut-off, shown next to final map) cryoEM map of bDENV-4 complexed with 3A06 IgG3. **(D)** Workflow of bZIKV complexed with 3A06 Fab displaying initial micrograph number, initial particle picks, multiple rounds of 2D classification and *ab initio* reconstruction of virion map. An in-house virion model of chimeric Powassan virus (bPOWV) was used as the initial model for 3D refinement resulting in a 4.6 Å (0.143 FSC cut-off, shown next to final map) cryoEM map. Symmetry expansion of the ASU was then performed as above followed by manual refinement on cisTEM resulting in a 3.1 Å (0.143 FSC cut-off) cryoEM map of the ASU.

```
CLUSTAL format alignment by MAFFT (v7.511)


ZIKV-Natal    IRCIGVSNRDFVEGMSGGTWVDVVLEHGGCVTVMAQDKPTVDIELVTTTVSNMAEVRSYC
DENV2-ET-300  MRCIGISNRDFVEGVSGGSWVDIVLEHGSCVTTMAKNKPTLDFELIKTEAKHPATLRKYC
DENV4-ET-288  MRCVGVGNRDFVEGVSGGAWVDLVLEHGGCVTTMAQGKPTLDFELTKTTAKEVALLRTYC
              :**:*:.*******:***:***:*****.***.**:.***:*:** .* ... * :*.**

ZIKV-Natal    YEASISDMASDSRCPTQGEAYLDKQSDTQYVCKRTLVDRGWGNGCGLFGKGSLVTCAKFA
DENV2-ET-300  IEAKLTNTTTASRCPTQGEPSLNEEQDKRFVCKHSMVDRGWGNGCGLFGKGGIVTCAMFT
DENV4-ET-288  IEASISNITTATRCPTQGEPYLKEEQDQQYICRRDLVDRGWGNGCGLFGKGGVVTCAKFL
               **.::: :: :*******. *.::.* :::*:: :***************.:**** *

ZIKV-Natal    CSKKMTGKSIQPENLEYRIMLSVHGSQHSGMIVNDTGHETDENRAKVEITPNSPRAEATL
DENV2-ET-300  CKKNMEGKVVQPENLEYTIVITPHSGEE----NAVGNDTGKHGKEIKVTPQSSITEAEL
DENV4-ET-288  CSGKITGNLVQIENLEYTVVVTVHNGDT----HAVGNDTSNHGVTATITPRSPSVEVKL
              *. :: *: :* ***** :::: *..:      : .*::*.::     :**.*. .*. *

ZIKV-Natal    GGFGSLGLDCEPRTGLDFSDLYYLTMNNKHWLVHKEWFHDIPLPWHAGADTGTPHWNNKE
DENV2-ET-300  TGYGTVTMECSPRTGLDFNEMVLLQMENKAWLVHRQWFLDLPLPWLPGADTQGSNWIQKE
DENV4-ET-288  PDYGELTLDCEPRSGIDFNEMILMKMKKKTWLVHKQWFLDLPLPWTAGADTSEVHWNYKE
               .:* : ::*.**:*:**.::  : *::* ****::** *:**** .****   :*  **

ZIKV-Natal    ALVEFKDAHAKRQTVVVLGSQEGAVHTALAGALEAEMDGAKGRLSSGHLKCRLKMDKLRL
DENV2-ET-300  TLVTFKNPHAKKQDVVVLGSQEGAMHTALTGATEIQM-SSGNLLFTGHLKCRLRMDKLQL
DENV4-ET-288  RMVTFKVPHAKRQDVTVLGSQEGAMHSALAGATEVDS-GDGNHMFAGHLKCKVRMEKLRI
               :* ** .***:* *.********:*:**:** * :  .  . : :*****:::*:**::

ZIKV-Natal    KGVSYSLCTAAFTFTKIPAETLHGTVTVEVQYAGTDGPCKVPAQMAVDMQTLTPVGRLIT
DENV2-ET-300  KGMSYSMCTGKFKVVKEIAETQHGTIVIRVQYEGDGSPCKIPFEI-MDLEKRHVLGRLIT
DENV4-ET-288  KGMSYTMCSGKFSIDKEMAETQHGTTVVKVKYEGAGAPCKVPIEI-RDVNKEKVVGRIIS
              **:**::*:. *.. *  *** *** .:.*:* * ..***:* ::  *::.   :**:*:

ZIKV-Natal    ANPVITESTENSKMMLELDPPFGDSYIVIGVGEKKITHHWHRSGS
DENV2-ET-300  VNPIVTEK--DSPVNIEAEPPFGDSYIIIGVEPGQLKLSWFKKG-
DENV4-ET-288  STPFAENT--NSVTNIELEPPFGDSYIVIGVGDSALTLHWFRKG-
               .*.  :.  :*   :* :********:***    :.  *.:.*
```

**Figure S3. Sequence alignment of DENV2, DENV4 and ZIKV E-proteins that were used in the cryo-EM analysis.** 3A06 antibody footprints are marked as stars in magenta color for HC over brown outline and pink color for LC in red outline.

```
CLUSTAL format alignment by MAFFT (v7.511)


DENV1-WP74      MRCVGIGNRDFVEGLSGATWVDVVLEHGSCVTTMAKDKPTLDIELLKTEVTNPAVLRKLC
DENV2-16681     MRCIGMSNRDFVEGVSGGSWVDIVLEHGSCVTTMAKNKPTLDFELIKTEAKQPATLRKYC
DENV3-UNC3001   MRCVGIGNRDFVEGLSGATWVDVVLEHGGCVTTMAKNKPTLDIELQKTEATQLATLRKLC
DENV4-SL92      MRCVGVGNRDFVEGVSGGAWVDLVLEHGGCVTTMAQGKPTLDFELTKTTAKEVALLRTYC
ZIKV-PRVABC59   IRCIGVSNRDFVEGMSGGTWVDVVLEHGGCVTVMAQDKPTVDIELVTTTVSNMAEVRSYC
                :**:*:.*******:**.:***:*****.***.**:.***:*:** .* ..: * :*. *

DENV1-WP74      IEAKISNTTTDSRCPTQGEATLVEEQDTNFVCRRTFVDRGWGNGCGLFGKGSLITCAKFK
DENV2-16681     IEAKLTNTTTESRCPTQGEPSLNEEQDKRFVCKHSMVDRGWGNGCGLFGKGGIVTCAMFR
DENV3-UNC3001   IEGKITNITTDSRCPTQGEAVLPEEQDQNYVCKHTYVDRGWGNGCGLFGKGSLVTCAKFQ
DENV4-SL92      IEASISNITTATRCPTQGEPYLKEEQDQQYICRRDVVDRGWGNGCGLFGKGGVVTCAKFS
ZIKV-PRVABC59   YEASISDMASDSRCPTQGEAYLDKQSDTQYVCKRTLVDRGWGNGCGLFGKGSLVTCAKFA
                 *...::: :: :*******. * ::.* .::*::  ***************.::*** *

DENV1-WP74      CVTKLEGKIVQYENLKYSVIVTVHTGDQHQ-----VGNETTEHGTTATITPQAPTSEIQ
DENV2-16681     CKKNMEGKVVQPENLEYTIVITPHSGEEHA-----VGNDTGKHGKEIKITPQSSITEAE
DENV3-UNC3001   CLEPIEGKVVQYENLKYTVIITVHTGDQHQ-----VGNET--QGVTAEITPQASTTEAI
DENV4-SL92      CSGKITGNLVQIENLEYTVVVTVHNGDTHA-----VGNDTSNHGVTATITPRSPSVEVK
ZIKV-PRVABC59   CSKKMTGKSIQPENLEYRIMLSVH-GSQHSGMIVNDTGHETDENRAKVEITPNSPRAEAT
                *   : *: :* ***:* :::: * *. *       .*::*  :     ***.:.  *

DENV1-WP74      LTDYGALTLDCSPRTGLDFNEMVLLTMEKKSWLVHKQWFLDLPLPWTSGASTSQETWNRQ
DENV2-16681     LTGYGTVTMECSPRTGLDFNEMVLLQMENKAWLVHRQWFLDLPLPWLPGADTQGSNWIQK
DENV3-UNC3001   LPEYGTLGLECSPRTGLDFNEMILLTMKNKAWMVHRQWFFDLPLPWTSGATTETPTWNRK
DENV4-SL92      LPDYGELTLDCEPRSGIDFNEMILMKMKKKTWLVHKQWFLDLPLPWTAGADTSEVHWNYK
ZIKV-PRVABC59   LGGFGSLGLDCEPRTGLDFSDLYYLTMNNKHWLVHKEWFHDIPLPWHAGADTGTPHWNNK
                *  :* : ::*.**:*:**.::  : *::* *:**::** *:**** .** *    *  :

DENV1-WP74      DLLVTFKTAHAKKQEVVVLGSQEGAMHTALTGATEIQTSGTT-TIFAGHLKCRLKMDKLT
DENV2-16681     ETLVTFKNPHAKKQDVVVLGSQEGAMHTALTGATEIQMSSGN-LLFTGHLKCRLRMDKLQ
DENV3-UNC3001   ELLVTFKNAHAKKQEVVVLGSQEGAMHTALTGATEIQNSGGT-SIFAGHLKCRLKMDKLE
DENV4-SL92      ERMVTFKVPHAKRQDVTVLGSQEGAMHSALAGATEVDSGDGN-HMFAGHLKCKVRMEKLR
ZIKV-PRVABC59   EALVEFKDAHAKRQTVVVLGSQEGAVHTALAGALEAEMDGAKGRLSSGHLKCRLKMDKLR
                : :* ** .***:* *.********:*:**:** * : .. .  : :*****:::*:**

DENV1-WP74      LKGMSYVMCTGSFKLEKEVAETQHGTVLVQVKYEGTDAPCKIPFSSQ-DEKGVTQNGRLI
DENV2-16681     LKGMSYSMCTGKFKVVKEIAETQHGTIVIRVQYEGDGSPCKIPFEIM-DLEKRHVLGRLI
DENV3-UNC3001   LKGMSYAMCTNTFVLKKEVSETQHGTILIKVEYKGEDAPCKIPFSTE-DGQGKAHNGRLI
DENV4-SL92      IKGMSYTMCSGKFSIDKEMAETQHGTTVVKVKYEGAGAPCKVPIEIR-DVNKEKVVGRVI
ZIKV-PRVABC59   LKGVSYSLCTAAFTFTKIPAETLHGTVTVEVQYAGTDGPCKVPAQMAVDMQTLTPVGRLI
                :**:** :*:  * . *  :** ***  :.*:* * ..***:* .   * :     **:*

DENV1-WP74      TANPIVTD--KEKPVNIEAEPPFGESYIVVGAGEKALKLSWFKKG-
DENV2-16681     TVNPIVTE--KDSPVNIEAEPPFGDSYIIIGVEPGQLKLNWFKKG-
DENV3-UNC3001   TANPVVTK--KEEPVNIEAEPPFGESNIVIGIGDNALKINWYKKG-
DENV4-SL92      SSTPLAEN--TNSVTNIELEPPFGDSYIVIGVGNSALTLHWFRKG-
ZIKV-PRVABC59   TANPVITESTENSKMMLELDPPFGDSYIVIGVGEKKITHHWHRSGS
                : .*:  .   :.   :* :****:* *::*     :.  *.:.*
```

**Figure S4. Sequence alignment of all four DENV serotypes and ZIKV strain used in the neutralization assays.** 3A06 antibody footprints are marked as stars in magenta color for HC over brown outline and pink color for LC in red outline.

```
CLUSTAL format alignment by MAFFT (v7.511)


conDENV1.v3     MRCVGIGSRDFVEGLSGATWVDVVLEHGKCVTVMMKDKPTLDIELLKTEVTNPAVLRKLC
conDENV2.v3     MRCIGISNRDFVEGVSGGSWVDIVLEHGKCVTVMMKNKPTLDFELIKTEAKQPATLRKYC
conDENV3.v3     MRCVGVGNRDFVEGLSGATWVDVVLEHGKCVTVMMKNKPTLDIELQKTEATQLATLRKLC
conDENV4.v3     MRCVGVGNRDFVEGVSGGAWVDLVLEHGKCVTVMMQGKPTLDFELTKTTAKEVALLRTYC
conZIKV.v3      IRCIGVSNRDFVEGMSGGTWVDVVLEHGKCVTVMMQDKPTVDIELVTTTVSNMAEVRSYC
                :**:*:..******:**.:***:************:.***:*:** .* ..: * :*. *

conDENV1.v3     IEAKISNTTTDSRCPTQGEATLVEEQDANFVCRRTFVDRGWGNGCDLFGKGSLITCAKFK
conDENV2.v3     IEAKLTNTTTASRCPTQGEPSLNEEQDKRFVCKHSMVDRGWGNGCDLFGKGGIVTCAMFT
conDENV3.v3     IEGKITNITTDSRCPTQGEAVLPEEQDQNYVCKHTYVDRGWGNGCDLFGKGSLVTCAKFQ
conDENV4.v3     IEASISNITTATRCPTQGEPYLKEEQDQQYICRRDVVDRGWGNGCDLFGKGGVVTCAKFS
conZIKV.v3      YEASISDMASDSRCPTQGEAYLDKQSDTQYVCKRTLVDRGWGNGCDLFGKGSLVTCAKFA
                 *..::: :: :*******. * ::.* .::*::  ***************.::*** *

conDENV1.v3     CVTKLEGKIVQYENLKYSVIVTVHTGDQHQ------VGNESTEHGTTATITPQAPTSEIQ
conDENV2.v3     CKKNMEGKIVQPENLEYTIVITPHSGEENA------VGNDTGKHGKEIKVTPQSSITEAE
conDENV3.v3     CLEPIEGKVVQYENLKYTVIITVHTGDQHQ------VGNET--QGVTAEITPQASTTEAI
conDENV4.v3     CSGKITGNLVQIENLEYTVVVTVHNGDTHA------VGNDTSNHGVTATITPRSPSVEVK
conZIKV.v3      CSKKMTGKSIQPENLEYRIMLSVH-GSQHSGMIVNDTGHETDENRAKVEITPNSPRAEAT
                *   : *: :* ***:* :::: * *. :       .*:::  :     :**.:.  *

conDENV1.v3     LTDYGALTLDCSPRTGLDFNEMVLLTMKEKSWLVHKQWFLDLPLPWTSGASTSQETWNRQ
conDENV2.v3     LTGYGTVTMECSPRTGLDFNEMVLLQMENKAWLVHRQWFLDLPLPWLPGADTQGSNWIQK
conDENV3.v3     LPEYGTLGLECSPRTGLDFNEMILLTMKNKAWMVHRQWFFDLPLPWTSGATTETPTWNRK
conDENV4.v3     LPDYGELTLDCEPRSGIDFNEMILMKMKKKTWLVHKQWFLDLPLPWTAGADTSEVHWNYK
conZIKV.v3      LGGFGSLGLDCEPRTGLDFSDLYYLTMNNKHWLVHKEWFHDIPLPWHAGADTGTPHWNNK
                *  :* : ::*.**:*:**.::  : *::* *:**::** *:**** .** *    *  :

conDENV1.v3     DLLVTFKTAHAKKQEVVVLGSQEGCMHTALTGATEIQTSGTT-TIWPGHLKCRLKMDKLT
conDENV2.v3     ETLVTFKNPHAKKQDVVVLGSQEGCMHTALTGATEIQMSSGN-LLWPGHLKCRLRMDKLQ
conDENV3.v3     ELLVTFKNAHAKKQEVVVLGSQEGCMHTALTGATEIQNSGGT-SIWPGHLKCRLKMDKLE
conDENV4.v3     ERMVTFKVPHAKRQDVTVLGSQEGCMHSALAGATEVDSGDGN-HMWPGHLKCKVRMEKLR
conZIKV.v3      EALVEFKDAHAKRQTVVVLGSQEGCVHTALAGALEAEMDGAKGRLWPGHLKCRLKMDKLR
                : :* ** .***:* *.********:*:**:** * : .. .  :*******:::*:**

conDENV1.v3     LKGMSYVMCTGSFKLEKEVAETQHGTVLVQIKYEGTDAPCKIPFSTQ-DEKGVTQNGRLI
conDENV2.v3     LKGMSYSMCTGKFKVVKEIAETQHGTIVIRVQYEGDGSPCKIPFEIM-DLEKRHVLGRLI
conDENV3.v3     LKGMSYAMCTNTFVLKKEVAETQHGTILIKVEYKGEDAPCKIPFSTE-DGQGKAHNGRLI
conDENV4.v3     IKGMSYTMCSGKFSIDKEMAETQHGTTVVKVKYEGAGAPCKVPIEIR-DVNKEKVVGRII
conZIKV.v3      LKGVSYSLCTAAFTFTKIPAETLHGTVTVEVQYAGTDGPCKVPAQMAVDMQTLTPVGRLI
                :**:** :*:  * . *  *** ***  :.::* * ..***:* .   * :     **:*

conDENV1.v3     TANPIVTD--KEKPVNIEAEPPFGESYIVIGAGEKALKLSWFKKG-
conDENV2.v3     TVNPIVTE--KDSPVNIEAEPPFGDSYIIIGVEPGQLKLSWFKKG-
conDENV3.v3     TANPVVTK--KEEPVNIEAEPPFGESNIVIGIGDNALKINWYKKG-
conDENV4.v3     SSTPFAEN--TNSVTNIELEPPFGDSYIVIGVGNSALTLHWFRKG-
conZIKV.v3      TANPVITESTENSKMMLELDPPFGDSYIVIGVGEKKITHHWHRSGS
                : .*.  .   :.   :* :****:* *:**     :.  *.:.*
```

**Figure S5. Sequence alignment of all four DENV serotypes and ZIKV recombinant E-dimer protein used in the binding and affinity assays.** 3A06 antibody footprints are marked as stars in magenta color for HC over brown outline and pink color for LC in red outline.

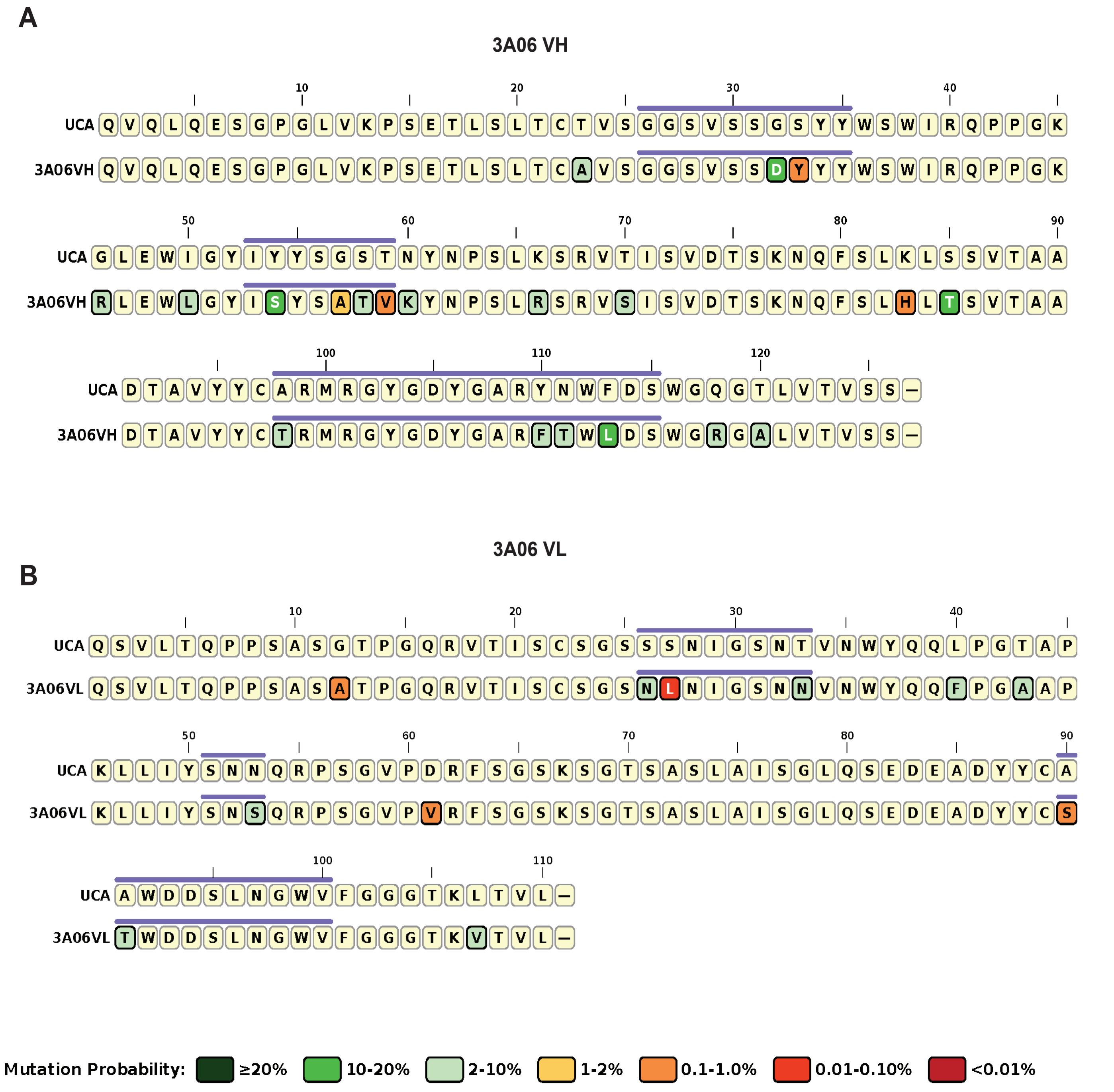


**Figure S6: ARMADiLLO server based improbable mutation analysis of 3A06 variable heavy and light chain genes.** Shown are the **(A)** heavy chain and **(B)** light chain sequence alignments with their respective germline UCA sequences.

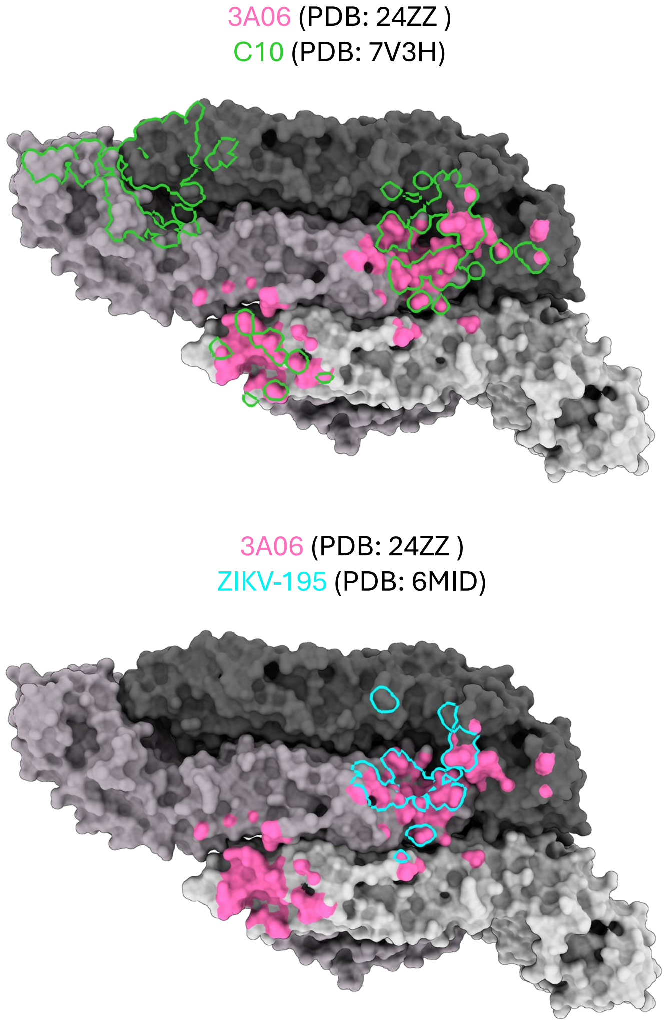


**Figure S7: Comparison of 3A06 footprints with existing cryo-EM structures of quaternary epitope targeting DENV or ZIKV mAbs.** Footprints of 3A06 depicted in pink surface coloring. Outline of comparator mAbs C10 (PDB: 7V3H), SIgN-3C (PDB: 7BU8), ZIKV-195 (6MID), and ADI30056 (PDB: 7KCR) overlayed onto 3A06 footprint in green, orange, blue and yellow, respectively.

**A**

**3A06 VH**

```
                                               FR1-IMGT          CDR1-IMGT       FR2-IMGT     CD
                                                (1-26)            (27-38)         (39-55)      (
                                    __________________________ ____________ _________________ __
                                    1        10        20        30         40        50
                                    .........|.........|...... ...|........ .|.........|..... ..
3A06_VH                             QVQLQESGP.GLVKPSETLSLTCAVS GGSVS..SDYYY WSWIRQPPGKRLEWLGY IS
M29811 Homsap IGHV4-61*01 F          QVQLQESGP.GLVKPSETLSLTCTVS GGSVS..SGSYY WSWIRQPPGKGLEWIGY IY
                                                           A           DY             R   L    S

                                    R2-IMGT                 FR3-IMGT
                                    56-65)                  (66-104)
                                    ________ _______________________________________
                                     60         70        80        90        100
                                    ..|..... ....|.........|.........|.........|....
3A06_VH                             YS...ATV KYNPSLR.SRVSISVDTSKNQFSLHLTSVTAADTAVYYC TR
M29811 Homsap IGHV4-61*01 F          YS...GST NYNPSLK.SRVTISVDTSKNQFSLKLSSVTAADTAVYYC AR
                                         ATV K     R    S            H T             T
```

**B**

**3A06 VL**

```
                                               FR1-IMGT          CDR1-IMGT       FR2-IMGT     CD
                                                (1-26)            (27-38)         (39-55)      (
                                    __________________________ ____________ _________________ __
                                    1        10        20        30         40        50
                                    .........|.........|...... ...|........ .|.........|..... ..
3A06_VL                             QSVLTQPPS.ASATPGQRVTISCSGS NLNI....GSNN VNWYQQFPGAAPKLLIY SN
AC279207 Homsap IGLV1-44*01 F        QSVLTQPPS.ASGTPGQRVTISCSGS SSNI....GSNT VNWYQQLPGTAPKLLIY SN
                                                A              NL         N       F  A

                                    R2-IMGT                 FR3-IMGT
                                    56-65)                  (66-104)
                                    ________ _______________________________________
                                     60         70        80        90        100
                                    ..|..... ....|.........|.........|.........|....
3A06_VL                             .......S QRPSGVP.VRFSGSK..SGTSASLAISGLQSEDEADYYC STWDDSLNG
AC279207 Homsap IGLV1-44*01 F        .......N QRPSGVP.DRFSGSK..SGTSASLAISGLQSEDEADYYC AAWDDSLNG
                                           S         V                               ST
```

**Figure S8: IMGT analysis generated alignment of 3A06 variable region amino acid sequences with their respective germline sequences.** Shown are the **(A)** heavy chain and **(B)** light chain sequence alignments.

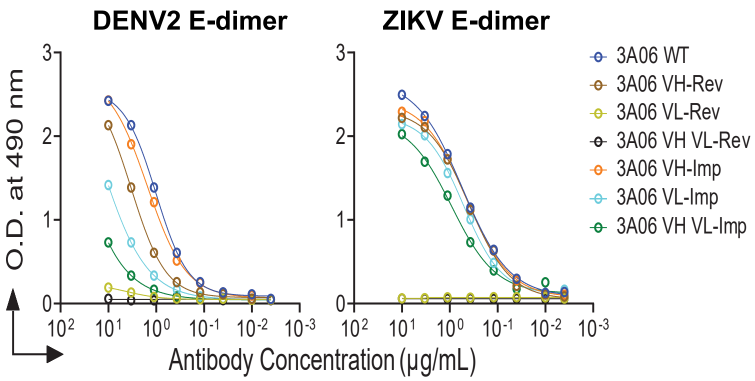


**Figure S9: Binding analysis of 3A06 VH or VL reverted variants.** Panel shows binding data of 3A06 heavy chain (VH) and light chain (VL) reverted variants, VH and VL improbable reverted variants to DENV2 and ZIKV E-dimer proteins.

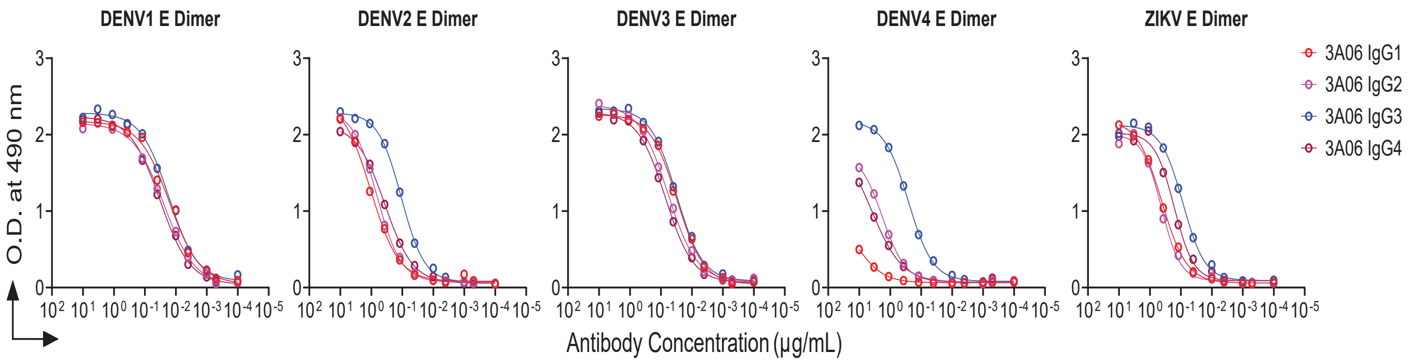


**Figure S10: 3A06 IgG subclasses binding analysis to DENV1-4 and ZIKV E-protein dimers.** Panel shows binding data of 3A06 IgG subclasses IgG1, IgG2, IgG3 and IgG4 to E-dimer proteins of all four DENV serotypes and ZIKV.

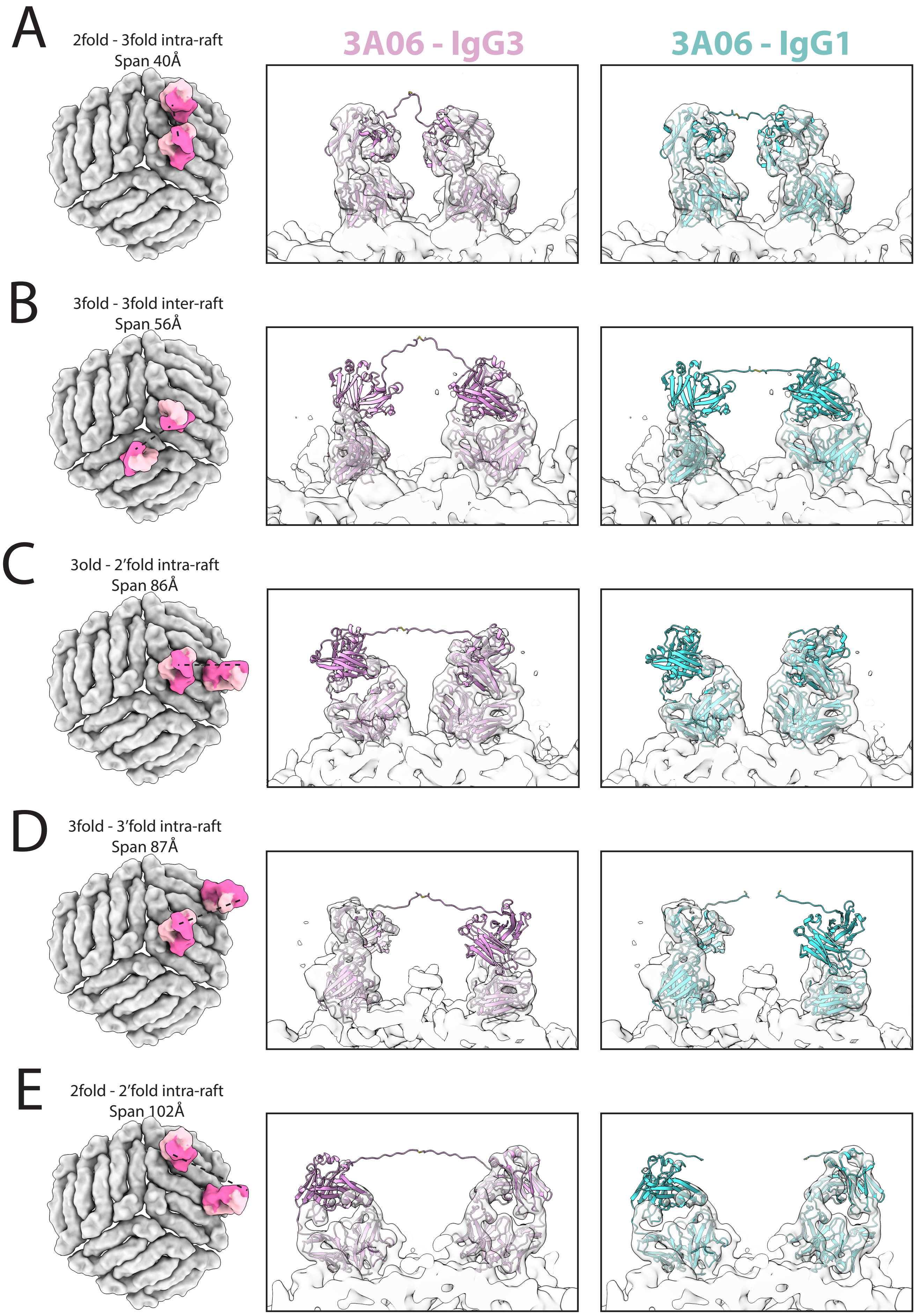

A
2fold - 3fold intra-raft
Span 40Å
3A06 - IgG3
3A06 - IgG1
B
3fold - 3fold inter-raft
Span 56Å
C
3old - 2'fold intra-raft
Span 86Å
D
3fold - 3'fold intra-raft
Span 87Å
E
2fold - 2'fold intra-raft
Span 102Å

**Figure S11: Comparison of bivalent engagement modes for 3A06 in both IgG3 and IgG1 format.** 3 rafts are shown for each mode (grey surface) with Fab densities shown with HC (magenta) and LC (pink). Right hand panels show cryoEM map of DENV2 and 3A06 IgG3 (white) fitted with either 3A06 on IgG3 framework (pink) or IgG1 (cyan). HC linker regions were modelled to test the compatibility of the relative linker regions with the HC position and epitope spacing **(A)** Intra-raft 2 to 3 -fold bivalent interaction results in a 40 Å span between HC Glu216 position (Eu numbering). **(B)** Inter-raft 3 to 3-fold site engagement results in a 56 Å span. **(C)** Inter-raft 3 to alternate 2 fold position results in a span of 86 Å. **(D)** Intra-raft 3 to 3-fold result in a span of 87 Å. **(E)** Intra-raft 2 to 2-fold results in a span of 102 Å.

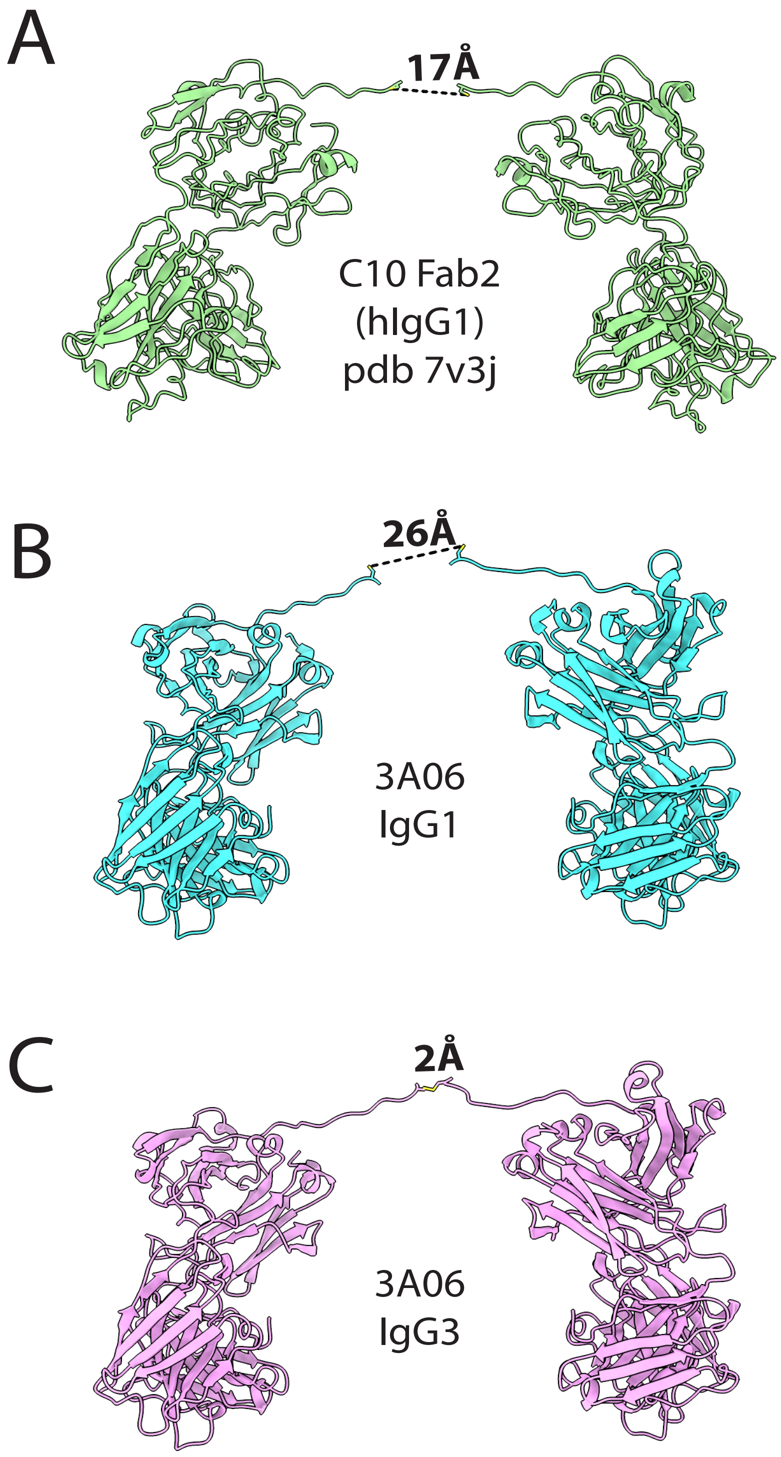


**Figure S12: Imposed HC hinge disulfide distances assuming simultaneous engagement of both 3-fold epitopes within a raft by a single bivalent antibody.** Distances measure between the Cys residue that forms the first HC hinge DS bridge (Cys226 for IgG1 and Cys229 for IgG3, Eu numbering). **(A)** The distance between Cys226 for IgG1 Fab2' of C10 is 17Å **(B)** The distance between Cys226 for 3A06 in IgG1 format is 27 Å **(C)** The distance between Cys229 for 3A06 in IgG3 format is 2 Å.

**Table S1: CryoEM data collection of bZIKV + 3A06 Fab.**

| | ***bZIKV + 3A06 Fab (PDB: 24ZZ, EMDB: EMD-69949)*** |
|---|---|
| ***Data collection and processing*** | |
| *Magnification* | 60,000x |
| *Voltage (kV)* | 300 |
| *Electron exposure (e-Å$^2$)* | 40 |
| *Number of frames per movie* | 40 |
| *Defocus range (μm)* | -2.5 to -0.5 |
| *Nominal Pixel size* | 0.40995 (Super-resolution) |
| ***Virion map*** | |
| *Symmetry Imposed* | I3 |
| *Initial particle number* | 8186 |
| *Final particle number* | 6483 |
| *Map Resolution (Å)* | 4.54 |
| *FSC threshold* | 0.143 |
| ***ASU map (symmetry expansion)*** | |
| *Symmetry Imposed* | C1 |
| *Final particle number* | 388,980 |
| *Map resolution (Å)* | 3.09 |
| *FSC threshold* | 0.143 |
| ***Refinement*** | |
| *Pixel Size (Å)* | 0.788 |
| *Initial model used* | Zika Virus (PDB: 6CO8) + Fab (AlphaFold3) |
| *Model resolution (Å)* | 3.1 (virus) |
| *FSC threshold* | 0.143 |
| ***Model composition*** | |
| *Non-hydrogen atoms* | 21170 |
| *Protein residues* | 2770 |
| *Ligands* | 4 |
| *Global CC (CCvol_)* | 0.87 |
| ***R.m.s deviations*** | |
| *Bond lengths (Å)* | 0.010 (0) |
| *Bond angles (°)* | 0.838 (3) |
| ***Validation*** | |
| *Clashscore* | 2 |
| *Poor rotamers (%)* | 0.04 |
| ***Ramachandran plot*** | |
| *Favoured (%)* | 95.16 |
| *Allowed (%)* | 4.84 |
| *Outliers (%)* | 0 |

### Table S2: CryoEM data collection.

| | bDENV-2 + 3A06 Fab (EMDB: 69863) | bDENV-2 + 3A06 IgG3 (EMDB: 69785) | bDENV-4 + 3A06 IgG3 (EMDB: 69800) |
|---|---|---|---|
| | | *Data collection and processing* | |
| **Magnification** | 60,000x | 60,000x | 60,000x |
| **Voltage (kV)** | 300 | 200 | 300 |
| **Electron exposure (e-Å²)** | 40 | 40 | 40 |
| **Number of frames per movie** | 40 | 40 | 40 |
| **Defocus range (μm)** | -2.5 to -0.5 | -2.5 to -0.5 | -2.5 to -0.5 |
| **Nominal Pixel size** | 0.40995 | 0.848 | 0.40995 |
| | | *Virion map* | |
| **Symmetry Imposed** | I3 | I3 | I3 |
| **Initial particle number** | 9,883 | 35,615 | 24,434 |
| **Final particle number** | 2,374 | 4,582 | 4,880 |
| **Map Resolution (Å)**<br>**FSC threshold** | 4.14<br>0.143 | 5.79<br>0.143 | 4.17<br>0.143 |
| | | *ASU map (symmetry expansion)* | |
| **Symmetry Imposed** | - | C1 | - |
| **Initial particle number** | - | 274,920 | - |
| **Final Particle Number** | - | *Class I*: 237,625<br>*Class II*: 240,646 | - |
| **Map Resolution (Å)** | - | *Class I*: 4.01<br>*Class II*: 3.24 | - |
| **FSC threshold** | | 0.143 | |
